# A chromosome-level genome assembly of the yellowfin seabream (*Acanthopagrus latus*) (Hottuyn, 1782) provides insights into its osmoregulation and sex reversal

**DOI:** 10.1101/2020.06.22.164046

**Authors:** Kecheng Zhu, Nan Zhang, Bao-Suo Liu, Liang Guo, Hua-Yang Guo, Shi-Gui Jiang, Dianchang Zhang

## Abstract

The yellowfin seabream *Acanthopagrus latus* is the economically most important Sparidae fish species in the northern South China Sea. As euryhaline fish, they are perfect model for investigating osmoregulatory mechanisms in teleosts. Moreover, the reproductive biology of hermaphrodites has long been intriguing; however, very little is known about the molecular pathways underlying their sex change. To elucidate genetic mechanisms of osmoregulation and sex reversal in this fish, a high-quality reference genome of the yellowfin seabream was generated by a combination of Illumina and PacBio technologies. The draft genome of yellowfin seabream was 806 Mb, with 732 Mb scaffolds anchored on 24 chromosomes. The contig N50 and scaffold N50 were 2.6 Mb and 30.17 Mb, respectively. The assembly is of high integrity and includes 92.23% universal single-copy orthologues based on benchmarking universal single-copy orthologs (BUSCO) analysis. Moreover, among the 19,631 protein-coding genes, we found that the *ARRDC3* and *GSTA* gene families related to osmoregulation underwent an extensive expansion in two euryhaline Sparidae fish genomes compared to other teleost genomes. Moreover, integrating sex-specific transcriptome analyses, several genes related to the transforming growth factor beta (TGF-β) signalling pathway involved in sex differentiation and development. This genomic resource will not only be valuable for studying the osmoregulatory mechanisms in estuarine fish and sex determination in hermaphrodite vertebrate species, but also provide useful genomic tools for facilitating breeding of the yellowfin seabream.

## 1. INTRODUCTION

Sparidae (order Spariformes) is a group of euryhaline fishes inhabiting the temperate to tropical waters of Atlantic, Indian and Pacific Oceans (Hesp et al., 2004). It is a family of teleosts with increased commercial significance, comprised approximately 38 genera and 159 species such as porgies and seabreams, some of which provide highly protein seafood (Natsidis et al., 2019; Tinacci et al., 2019). From the dataset of the International Union for Conservation of Nature’s Red List Process, a global extinction risk assessment of Sparidae showed that approximately 25 species are threatened or near-threatened according to their body weight (Comeros-Raynal et al., 2016). Moreover, Food and Agriculture Organization (FAO) also reported that the advantage of worldwide aquaculture production, contributing to 47% of total marine products, also underlined the over-fishing of Sparidae (FAO, 2018). The Sparidae has been widely favored for its high-quality protein and recognized important aquaculture species in China with annual yield accounting 81,107 tons in 2017 (FBMA, 2018). Therefore, changes in Sparidae species and their abundances can dramatically affect the survivability and health of this highly complex ecosystem.

The Sparidae is a typical euryhaline estuarine fish, living in a wide range of water salinities. Osmoregulation is one of the important factors that dominate the accommodative ability of an organism to response to different salinities and is a meaningful process for realizing the physiological function of commercial fishes to promote aquaculture situations and production (Jaffer et al., 2020). The regulatory mechanism of salinity stress was regulated by a plentiful of osmoregulatory effector genes and their protein products (Kültz 2015; Xu et al., 2015; Chen et al., 2019), which included Na^+^/K^+^/2Cl^−^ co-transporter (NKCC), cystic fibrosis transmembrane conductance regulator (CFTR), several plasma membrane ATPases, and other transporters (Marshall 2013). The osmosensory signalling networks, controlling above effector genes, are most obvious and effective in euryhaline species involving in osmoregulatory mechanism (Kültz 2012). Moreover, several genomic datasets of Sparidae fish have been published in recent years, such as those for *A. schlegelii* (Zhang et al., 2018), *S. aurata* (Pauletto et al., 2018), and red seabream (*Pagrus major*) (Shin et al., 2018). However, no genomic research has been conducted thus far to identify the candidate genes and pathways for survival over a large range of salinities, which severely hinders studies examining their osmoregulatory mechanisms. Sparidae is a euryhaline species with the capacity to cope with demands over a wide range of salinities and thus is a perfect model fish to study osmoregulatory responses to salinity-adaptive processes (Laiz-Carrión et al., 2005; Cook and Herbert 2012; Ruiz-Jarabo et al., 2018; Kisten et al., 2019). As such, understanding the osmoregulatory mechanisms and developing genomic selection programmes have become important goals of Sparidae research.

The Sparidae has a striking life cycle with a sequential protandrous hermaphrodite physiological phenomenon. It matures first as male and in the following cycles-depending on social factors, growth, and diet-female reproductive organs develop. Numerous genes possessed higher expression in one sex than the other, and sex-biased transcription is identified to be the manner by which sexual dimorphisms can be resulted from the genome (Parsch and Ellegren, 2013). Furthermore, the particular genetic mechanism of sex determination (SD) genes is elucidated in a small number of species, such as *Dmrt1bY*/*DmY* in medaka species *Oryzias curvinotus* and *Oryzias latipes* (Matsuda et al., 2002), anti-Müllerian hormone (*Amhy*) in *Odontesthes hatcheri* (Patagonian pejerrey) (Hattori et al., 2012), anti-mullerian hormone receptor 2 (*Amhr2*) in Fugu species *Takifugu poecilonotus, Takifugu rubripes*, and *Takifugu pardalis* (Kamiya et al., 2012), gonadal soma derived factor (*Gsdf*) in medaka species *Oryzias luzonensis* (Myosho et al., 2012), sexually dimorphic on the Y-chromosome (*SdY*) in *Oncorhynchus mykiss* (rainbow trout) (Yano et al., 2012), breast cancer anti-resistance 1 (*BCAR1*) in *Ictalurus punctatus* (channel catfish) (Bao et al., 2019). Moreover, sex-determining regions (sex chromosome) or sex-linked markers (SNP) have already been reported in some fish, such as *Cynoglossus semilaevis* (Half-smooth tongue sole) (Shao et al., 2010; Chen et al., 2014; Cui et al., 2017), fugu Takifugu rubripes (Kamiya et al., 2012), and *Dicentrarchus labrax* (European sea bass) (Palaiokostas et al., 2015), and so on. Additionally, in gilthead sea bream (*Sparus aurata*), fast evolution is observed only for highly ovary-biased genes due to female-specific patterns of selection that are related to the peculiar reproduction mode (Pauletto et al., 2018). By performing a comparative genomic analysis, Zhang et al. (2018) found three types of genes that are potentially associated with sex reversal and are useful for studying the genetic basis of protandrous hermaphroditism in black seabream (*Acanthopagrus schlegelii*). Moreover, comparative transcriptomic analyse has been conducted to identify the male- and female-specific genes and pathways that are most likely involved in sex maintenance in Sparidae (Tsakogiannis et al., 2019). Although above research has been performed in Sparidae, the genetic mechanism of sex change is still not fully understood. Consequently, to elucidate their interesting biological characteristics, the yellowfin seabream (*Acanthopagrus latus*) is also a good model for investigating the genetic mechanisms of sex reversal.

As one of the most popular and valuable commercial marine fishes in China and Indo-Western Pacific Ocean, the *A. latus* (Fig. 1), has some interesting characteristics such as high economic value, good meat quality, resistance to diseases, and good adaptability to various environments (Bauchot & Smith, 1986; Carpenter, 2001). The osmoregulatory processes are accomplished by variety of transporters and enzymes, and the synthesis and responding of these proteins are extremely energy consuming in Sparidae. However, the regulatory mechanisms enabling osmoregulation remain poorly understood for this economically important species. Moreover, studies in fish have already redefined our understanding of the complexity and plasticity of the sex determination (SD) and sex reversal process, yet little is known about the detailed genetic information underlying sex change. Here, a high-quality genome assembly is necessary to understand the functional, ecological and evolutional genomics of this species and of other Sparidae. We present a chromosome-level genome assembly and a draft assembly of *A. latus*. The chromosome-level genome assembly and annotations are helpful for understanding the molecular basis of sex reversal and the characteristics of the osmoregulation. The genomic resources of these two assemblies can also facilitate future investigations of the evolution and biology of this species and provide a valuable resource for conservation and breeding management of *A. latus*.

**Figure 1.**
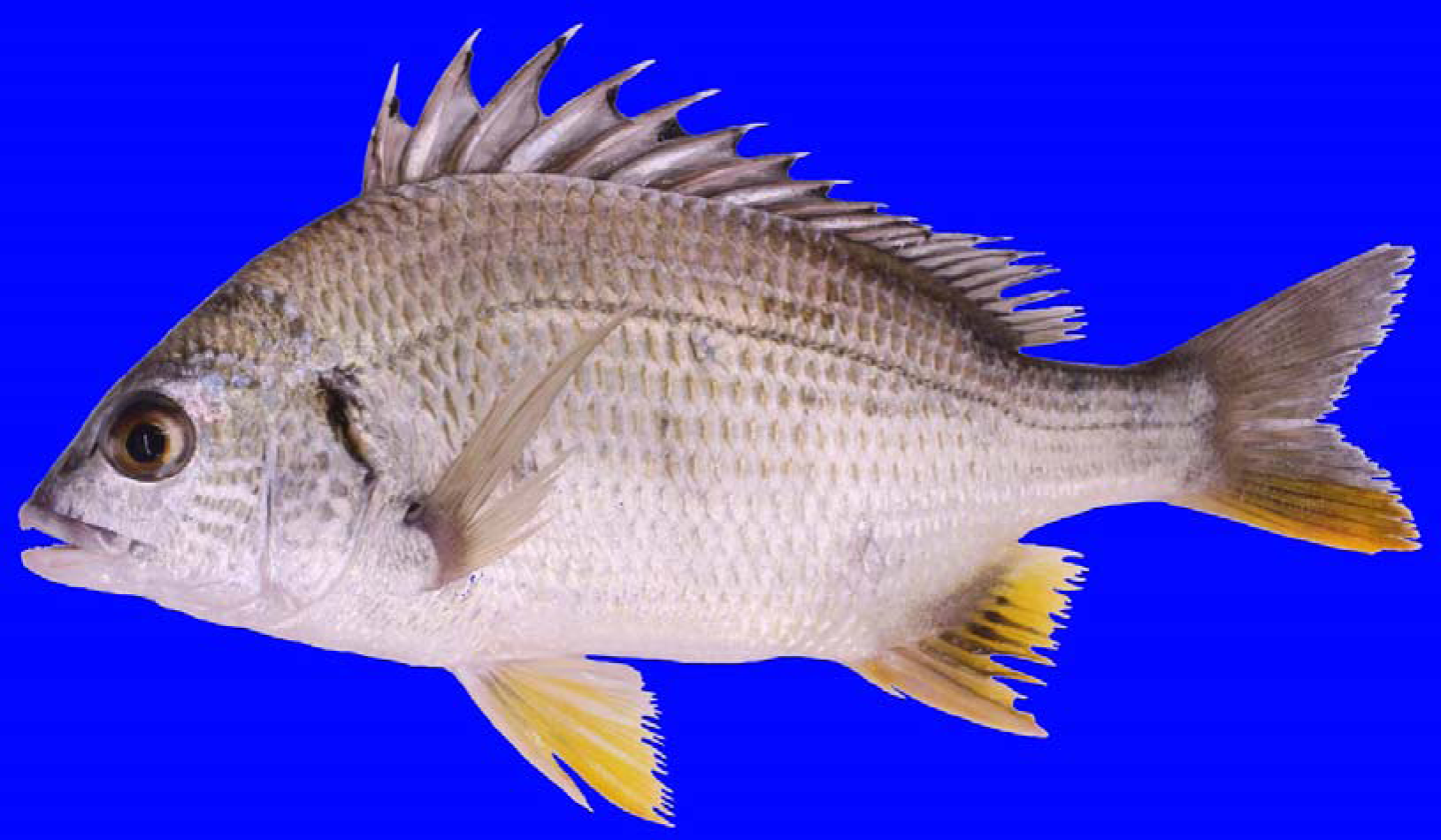
Image of an adult yellowfin seabream (*Acanthopagrus latus*).

## 2. MATERIALS AND METHODS

### 2.1 Ethics statement

All fish sampling in this study was permitted by the Animal Care and Use Committee of South China Sea Fisheries Research Institute, Chinese Academy of Fishery Sciences (No. SCSFRI96-253), and conformed with the regulations and guidelines established by this committee.

### 2.2 Sampling and sequencing

Female *A. latus* individual was obtained from a culture population in Yangjiang, Guangdong, China. Total genomic DNA was extracted from the muscle of female fish using a DNA extraction kit (Magen, Guangzhou, China) following the manufacturer’s protocols. The quality and quantity (concentration) of total DNA were determined by 1% agarose gels and a NanoDrop 2000 spectrophotometer (Thermo Scientific, USA), respectively.

Two short-insert size (180 and 500 bp) libraries and four long-insert size (3K, 5K, 10K and 14K bp) libraries were constructed according to standard Illumina procedures (Illumina, San Diego, CA, USA). The libraries were sequenced on a HiSeq 2500 system with the 250 bp PE or 150 bp PE modes (Table 1). Raw data were generated by an Illumina HiSeq 2000 sequencer and an Illumina Genome Analyzer IIx sequencer. Furthermore, two 20 kb libraries were also constructed by the extracted DNA samples following the PacBio manufacturing protocols (Pacific Biosciences, CA, USA). Subsequently, the libraries were sequenced with two cells based on the PacBio Sequel platform (Table 1).

**Table 1.**
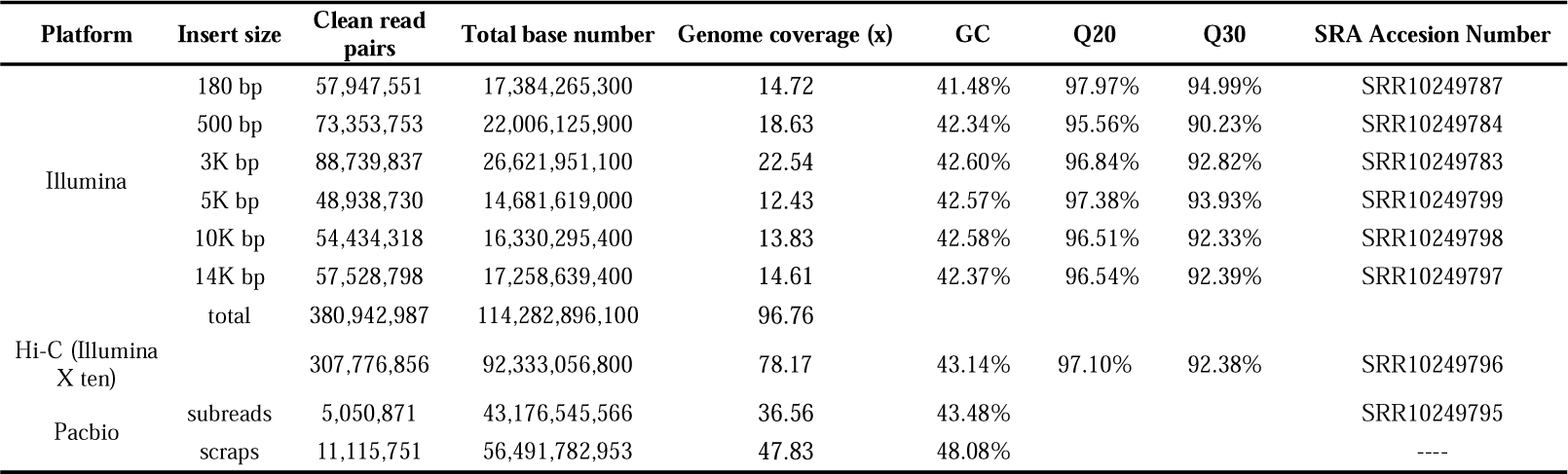
Data statistics of whole genome sequencing reads of female yellow sea bream.

| Platform | Insert size | Clean read pairs | Total base number | Genome coverage (x) | GC | Q20 | Q30 | SRA Accession Number |
| --- | --- | --- | --- | --- | --- | --- | --- | --- |
| Illumina | 180 bp | 57,947,551 | 17,384,265,300 | 14.72 | 41.48% | 97.97% | 94.99% | SRR10249787 |
|  | 500 bp | 73,353,753 | 22,006,125,900 | 18.63 | 42.34% | 95.56% | 90.23% | SRR10249784 |
|  | 3K bp | 88,739,837 | 26,621,951,100 | 22.54 | 42.60% | 96.84% | 92.82% | SRR10249783 |
|  | 5K bp | 48,938,730 | 14,681,619,000 | 12.43 | 42.57% | 97.38% | 93.93% | SRR10249799 |
|  | 10K bp | 54,434,318 | 16,330,295,400 | 13.83 | 42.58% | 96.51% | 92.33% | SRR10249798 |
|  | 14K bp | 57,528,798 | 17,258,639,400 | 14.61 | 42.37% | 96.54% | 92.39% | SRR10249797 |
|  | total | 380,942,987 | 114,282,896,100 | 96.76 |  |  |  |  |
| Hi-C (Illumina X ten) |  | 307,776,856 | 92,333,056,800 | 78.17 | 43.14% | 97.10% | 92.38% | SRR10249796 |
| Pacbio | subreads | 5,050,871 | 43,176,545,566 | 36.56 | 43.48% |  |  | SRR10249795 |
|  | scraps | 11,115,751 | 56,491,782,953 | 47.83 | 48.08% |  |  | ---- |

The Hi-C library sequencing technique was used to construct chromosome-level assemblies (Dudchenko et al., 2017; Shao et al., 2018; Zhou et al., 2019). Blood samples were fixed with formaldehyde, and a restriction enzyme (Mbo *I*) was added to digest the DNA, followed by repairing the 5’ overhang using a biotinylated residue (Rao et al., 2014). The fragments close to each other in the nucleus during fixation were ligated. A PE library with an approximately 300 bp insert size was constructed following the Hi-C library protocol (Shao et al., 2018). The library was sequenced on the Illumina HiSeq X Ten platform with in the 150 bp PE mode.

The reads from the Hi-C library sequencing were mapped to the polished genome using Bowtie2 (Langmead and Salzberg, 2012) with default parameters. Traditional read mapping led to a low mapping ratio because most of the reads were chimaeras, according to the Hi-C library construction procedures. To overcome this problem, an iterative alignment strategy was applied to improve the mapping ratio (Imakaev et al., 2012). Although the sequencing data were paired end, we aligned both reads independently. The read pairs with both ends uniquely mapped to the genome were collected for analysis. As in previous methods (Imakaev et al., 2012; Servant et al., 2015), the hiclib Python library (Imakaev et al., 2012) was applied to evaluate the interaction strengths between whole-genome contig sequences. Based on the interaction matrix, Lachesis (Burton et al., 2013) was used to anchor the contigs to the chromosomes using an agglomerative hierarchical clustering method.

To facilitate gene prediction, thirteen types of tissues (e.g., white muscle, stomach, gill, liver, kidney, spleen, intestine, fin, heart, eye, skin, brain, and male, female, and hermaphrodite gonads) were collected from male, female, and hermaphrodite individuals for full-length transcriptome sequencing on the Illumina HiSeq 4000 platform based on single-molecule real-time (SMRT) sequencing technology.

### 2.3 Read filtration and genome size estimation and assembly

All genomic and transcriptomic sequencing reads were cleaned using SolexaQA (Cox et al. 2010) to filter the low-quality bases and short reads < 25 bp. Before genome assembly, we estimated genome size with the cleaned reads from the two paired-end Illumina libraries using Jellyfish v2.020 (Marcais & Kingsford, 2011). After obtaining K-mers from the short-insert size (<1 Kb) reads with a 1 bp sliding window, the frequency of each K-mer was calculated. The K-mer frequencies fit a Poisson distribution when sufficient amount of data are present. The total genome size was deduced from these data in the following manner: genome size = K-mer num/Peak depth. Moreover, all cleaned genomic reads were assembled using Platanus (Kajitani et al. 2014), and the gaps in the scaffolds were closed with reads from the paired-end libraries.

To anchor the scaffolds into chromosomes, HiCUP v0.6.126 was first used to map and process the reads from the Hi-C library. Read pairs were mapped to the polished scaffolds using Bowtie2 with default parameters (Langmead and Salzberg, 2012). If both reads from one pair were uniquely mapped to the assembly, this pair was retained for the downstream filtration. HiCUP removed invalid pairs generated from contiguous sequences, circularization, dangling ends, internal fragments, re-ligation, PCR duplication, and fragments of the wrong size.

The completeness of the assembled genome was evaluated using benchmarking universal single-copy orthologs (BUSCO) version 3.0 (Simão et al., 2015) with the actinopterygii_odb9 database.

### 2.4 Repetitive element annotation

Before predicting protein-coding genes, we masked the repetitive regions of the assembly using a combination of ab initio and homology-based approaches. We used TRF (v.4.09) (Benson, 1999), RepeatMasker (v4.0.7), and RepeatProteinMask (Tarailo-Graovac and Chen, 2009) to detect and classify different types of repetitive sequences by aligning the genome sequences against the Repbase library (v.17.01) (Bao et al., 2015). Novel repeats were predicted using RepeatModeler (version 1.05) based on the de novo repeat library (Maziade et al., 1996). By using RepeatMasker (v.3.3.0), the transposable elements (TEs) in the genome were identified. The consensus non-redundant library is obtained by a combination of known, novel and tandem repeats.

### 2.5 Homology-based prediction

Functional annotation of the protein-coding genes in *A. latus* was performed. The protein sequences of sixteen species were aligned to the *A. latus* assembly using BLAST (E-value ≤1e-5) with the SwissProt and TrEMBL subsets of the UniProt database (Boeckmann et al., 2003), and the matches with length coverages >30% of the homologous proteins were considered gene-model candidates. Then, the corresponding homologous genome sequences were aligned against the matching proteins by using GeneWise (Birney et al., 2014) to improve the gene models.

To retrieve the biological pathways related to the *A. latus* genome, KEGG orthologous gene information was obtained using the KEGG Automatic Annotation Server (KAAS) (Moriya et al., 2007) with a representative eukaryote set as a reference and with default parameters; we linked the genes to KEGG pathways using a custom Perl script. To investigate specific gene features of *A. latus*, functional annotation with fifteen other sequenced teleost genomes was performed using the same procedure mentioned above.

### 2.6 Phylogenetic tree reconstruction and divergence time estimation

To detect variations in the *A. latus* genome, the 367 single-copy gene families conserved among *S. aurata, A. schlegelii, Danio rerio* (zebrafish), *Gadus morhua* (cod), *Labrus bergylta* (ballan wrasse), *Lepisosteus oculatus* (spotted gar), *Oreochromis niloticus* (tilapia), *O. latipes* (medaka), *Poecilia formosa* (Amazon molly), *Takifugu rubripes* (takifugu), *Tetraodon nigroviridis* (tetraodon), *Xiphophorus maculatus* (platyfish), *Cyprinus carpio* (common carp), *Larimichthys crocea* (large yellow croaker), and *Notothenia coriiceps* (black rockcod) were extracted and concatenated into a single data set by the MUSCLE program (MUSCLE, RRID:SCR 011812) and PhyML (PhyML, RRID:SCR 014629) (Guindon and Gascuel, 2003; Edgar, 2004; Guindon et al., 2010). To reduce the error in the topology of the phylogeny due to alignment inaccuracies, we used Gblocks (codon model) to remove unreliably aligned sites and gaps in the alignments. The phylogenetic tree was constructed and divergence times were calculated using the PAML 3.0 package (Yang and Rannala, 2006; Yang, 2007).

### 2.7 Gene family expansion and contraction analysis

Gene family expansion and contraction analyses were performed by CAFE 3.0 (De Bie et al., 2006). We downloaded osmoregulation-related genes, such as members of the ADDRC3, GSTA and SLC12A families, from the Ensemble or Genebank database and predicted their candidates using BLAST with those from eight other genomes. GeneWise was used to determine copy numbers and pseudogenes produced by frame shifts, and these were removed. The phylogenetic analysis of the expanded gene families was based on maximum likelihood and performed in MEGA 6.0 (Tamura et al., 2013), and the phylogenetic tree was visualized by EvolView (He et al., 2016).

### 2.8 Functional analysis of osmoregulation-related genes under acute salinity stress

To study whether these three families (e.g., ADDRC3, GSTA and SLC12A) are involved in the response to salinity changes in yellowfin seabream, an acute salinity stress experiment was implemented. One hundred and twenty fish (30 ± 5 g) were randomly and evenly transferred simultaneously to 5‰ and 35‰ salinity conditions from normal salinity (20‰) conditions with three replicates in each group. We then observed and recorded the behaviours of juvenile fish under salinity stress, with the juveniles removed if they lost consciousness. Finally, for the osmoregulatory tissues (e.g., gills, kidney, skin, and brain), samples were taken at 0, 6, and 12 h from each group (3 fish/replicate/time point). The fish were sampled under anaesthesia (MS-222). The tissues were immediately stored in liquid nitrogen for later use. Total RNA was extracted from the tissues, and first-strand cDNA was synthesized from 2 μg of total RNA and used as a template for quantitative real-time PCR (qRT-PCR) which was performed according to Zhu et al (2019). The mRNA levels of the target genes were quantified using the 2^−ΔΔCt^ method (Livak and Schmittgen, 2001).

### 2.9 Transcriptome analyses of the developmental process of gonads

To better comprehend the morphological types of gonads in different periods for the yellowfin seabream, the gonads were collected and fixed overnight in 4% paraformaldehyde. For sectioning, the fixed tissues were dehydrated using a graded series of ethanol concentrations (70–100%), followed by embedding in paraplast (Leica, Germany). Transverse sections for histological analyses were cut at 5 µm intervals on a retracting microtome and stained with haematoxylin and eosin (HE), following the method described by Chen et al. (2015).

*A. latus* specimens belonging to three different developmental stages of the gonads (stage 1: male; stage 2: intersexual; and stage 3: female) were used. RNA was extracted from gonad samples at the different stages of *A. latus* using RNAisoPlus Reagent (TaKaRa, Japan) according to the manufacturer’s instructions. The integrity and purity of the RNA were determined by gel electrophoresis and a NanoDrop ND-1000 (Thermo Scientific), respectively, before preparing the libraries for sequencing. Paired-end RNA sequencing was performed using the Illumina HiSeq 2000 platform. Low-quality score reads were filtered out, and the clean data were aligned to the reference genome using Bowtie (Langmead and Salzberg, 2012). For details of the analysis method, please refer to Tsakogiannis et al. (2019).

## 3 RESULTS AND DISCUSSION

### 3.1 Genome size estimation and assembly

Before sequencing, the genome size of female individuals of *A. latus* was estimated by K-mer depth and frequency distribution (Fig. S1). We prepared genomic DNA from a cultured female adult and performed whole-genome shotgun sequencing on three next-generation sequencing platforms, namely, Roche 454, Illumina and SOLiD, using both single-end and paired-end or mate-pair libraries of various insert sizes ranging from 180 bp to 14 kb (Table 1, Fig. S2). The distributions of the insert sizes in six libraries indicated that the *A. latus* genome is a single genome. The genome size is estimated to be 875 Mb according to the frequency distribution of K-mers (Fig. S1). After data filtering, we generated 381 million reads with a total length of 85.99 Gb from the adult female (Table 1), which was shorter than the length estimated by K-mer depth and frequency distribution.

We assembled an 806 Mb reference genome sequence from 192.0 Gb (approximately 162.6-fold coverage) of clean data. The contig and scaffold N50 lengths reached 2.6 Mb and 30.7 Mb, respectively. Meanwhile, the scaffold N90 length reached 17.4 Mb, and 90% of the assemblies were composed of 4,267 scaffolds, which were all greater than 195.6 kb in length (Table 2). These reads yielded a draft genome covering 96.18% of the estimated genome size with remaining unclosed gaps of approximately 1.46 Mb (0.18% of the total scaffold sequence) (Table 2). In the genomes of Sparidae fish, similar genome sizes have been reported for two other species, namely, *S. aurata* (830 Mb) (Pauletto et al., 2018) and *P. major* (829 Mb) (Shin et al., 2018); however, the genome size of *A. schlegelii* was 688 Mb (Zhang et al., 2018), which was smaller than that of the above three Sparidae fish. Moreover, the draft assemblies of *A. latus, S. aurata*, and *P. major* produced similar N50 contig lengths (2.6 Mb, 3.6 Mb, and 2.8 Mb, respectively), despite the 17.2-kb-smaller genome of *A. schlegelii*. Nevertheless, an N50 scaffold length of 30.7 Mb has been reported for the *A. latus* genome, while the N50 scaffold lengths for *S. aurata* and *A. schlegelii* described here were only 3.7 Mb and 7.6 Mb, respectively (no data for *P. major*). These results suggested the highest assembly quality for the *A. latus* genome. As is the case in this study, only Illumina paired-end sequencing has been performed for the *P. major* genome. To increase scaffold length, the addition of longer reads from a second technology would be necessary.

**Table 2.** Summary of yellow sea bream genome assembly.

| **Genome assembly statistics** |  |
| --- | --- |
| Total genome size (Mb) | 806 Mb |
| Length of gaps (N Number) | 1,461,838 |
| Contig N50 length (bp) | 2,603,585 |
| Scaffold Number | 4,267 |
| Scaffold N50 length (bp) | 30,688,393 |
| Scaffold N90 length (bp) | 17,436,492 |
| **Genome characteristics** |  |
| GC content (%) | 42.07 |
| No. of predicted protein-coding genes | 19,631 |
| Content of transposable elements (%) | 18.98 |
| Content of SSRs (%) | 1.07 |
| Chromosome average length (bp) | 30,506,010 |
| Quantity of scaffolds anchored on linkage groups | 24 |
| Average gene length (bp) | 9,468 |
| Length of scaffolds anchored on LGs (Mb) | 732 Mb |

The assembly quality was evaluated using two metrics: (1) the insert size distributions of the paired-end/mate-pair libraries by aligning reads to the genome using BWA (Li and Durbin, 2009) and (2) the ratio of RNA-sequencing read mapped to the genome using HISAT2 (Kim et al., 2015). First, the insert size distributions of the six libraries were consistent with the estimated insert sizes (Fig. S2). Second, the unigenes from the transcriptomes of gonads from three periods of the *A. latus* genome and approximately 90.00% of the unigenes exhibited significant hits in the genome (Table S1) (Kim et al., 2015). These two metrics indicated the high quality of this assembly.

The completeness of *A. latus* genes was evaluated by using BUSCO software (Simao et al., 2015). To further estimate the completeness of the assembly and gene prediction, BUSCO (RRID: SCR 015008) was conducted, and the results showed that the assembly contained 92.23% complete and 33.6% single-copy vertebrate BUSCO orthologues (Table S2). These high mapping rates indicated that the genome assembly of *A. latus* had high coverage and quality.

### 3.2 Genome annotation

De novo prediction and homology searching against the Repbase database showed that the identified repeat sequences covered 18.98% of the assembled genome in *A. latus* (Table S3). DNA transposons (1.51%), long interspersed nuclear elements (LINEs, 1.95%), long terminal repeats (LTRs, 1.00%), short interspersed nucleotide elements (SINEs, 0.01%), and unclassified (14.50%) were the categories of repetitive elements in the *A. latus* genome. Interspecific comparisons showed that the frequencies of the DNA transposons, LINEs, LTRs and SINEs were lower in *A. latus* than in other Sparidae fish, such as *S. aurata* (Table S4) (Pauletto et al., 2018) and *A. schlegelii* (Table S5) (Zhang et al., 2018).

Based on the repeat-masked genome assembly, we integrated de novo, homology searching-based and transcript-based methods to predict a protein-coding gene set comprising 19,631 genes (Table 2). The average gene length was 9,468 bp, which was consistent with the distributions of gene features in other teleosts.

### 3.3 Chromosome sequence assembly and synteny analysis

In addition, chromosome genome assembly is pivotal for genome comparisons and evolutionary studies (Fan et al., 2019a; He et al., 2019). Using the Hi-C library and PacBio sequencing, we obtained ∼92.3 Gb and 43.1 Gb of clean reads for the Hi-C and PacBio analyses, respectively (Table 1). Subsequently, combining these two sequencing datasets and our mapping strategy after read-quality filtering, we found that 4,267 scaffolds comprised 90.8% (732 Mb) of the *A. latus* genome assembly to form 24 linkage groups (LGs) (Fig. 2A, Table 2, Table S6), which was consistent with the findings of previous karyotype analyses of *A. latus* (Liu et al., 1991). This study provided the first chromosome-level genome of the Sparidae and provided a valuable genomic resource for evolutionary studies of *A. latus* and related species. However, there were still 128 unanchored sequences after PacBio-based chromosome construction with an N50 length of 53.7 kb, which was significantly smaller than that of the anchored sequences (5.9 Mb, 0.81% of the total anchored sequences) (Fig. 2A). The above length analysis and statistics demonstrated that continuity at the scaffold level not only delivered more sequential information but also may have influenced the assembly results during the PacBio-based chromosome construction process.

**Figure 2.**
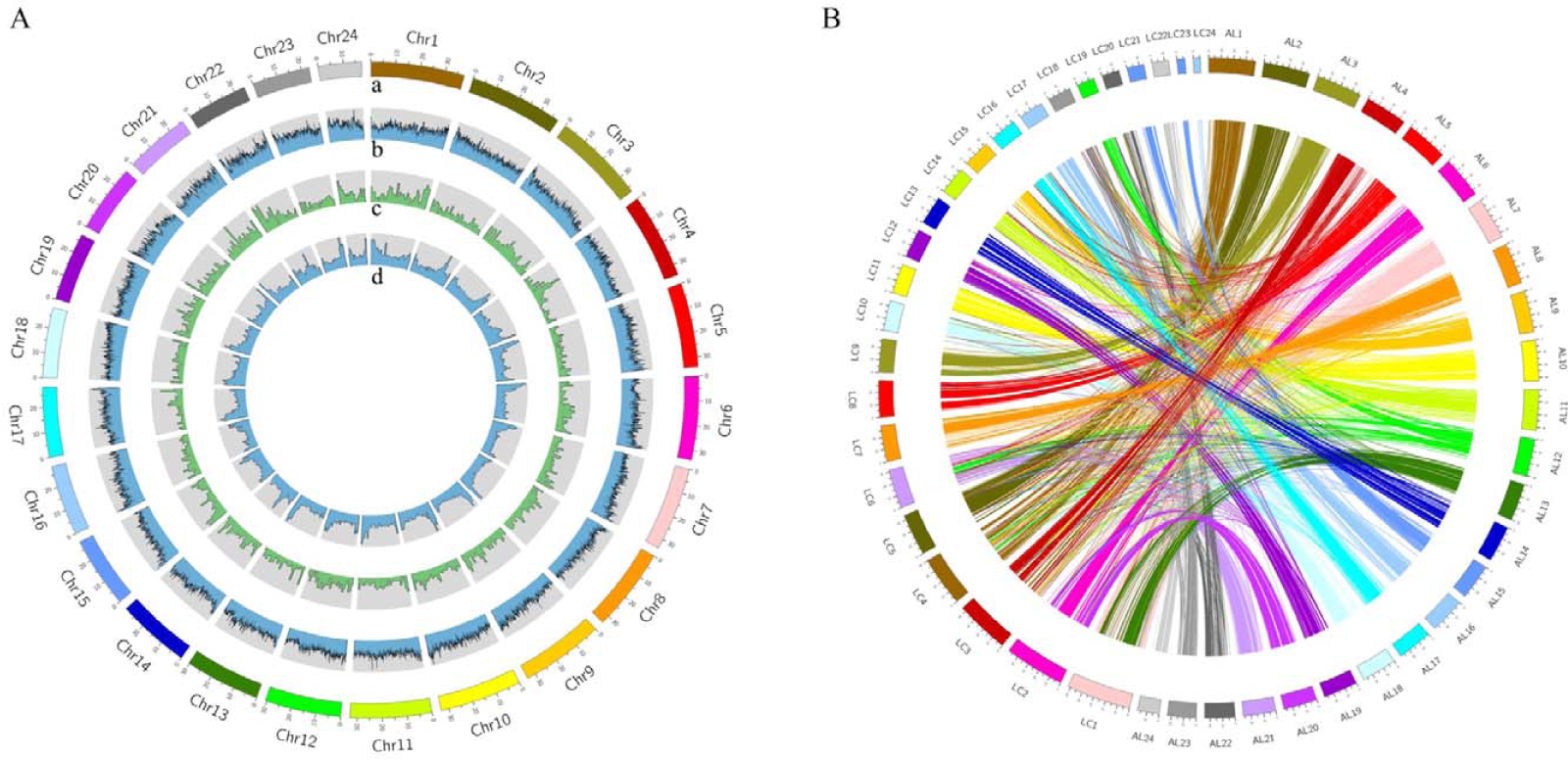
Global view of the *A. latus* genome and syntenic relationship between *A. latus* and *L. crocea*. (A) Global view of the *A. latus* genome. From outside to inside, 24 linkage groups (LGs) (a); GC content with sliding windows of 50-kb (b); gene distribution on each chromosome with sliding windows of 1 Mb (c); repeat content (d). (B) Syntenic relationship between the *A. latus* and *L. crocea* chromosomes.

A subsequent comparison of gene order between *A. latus* and its close relative *L. crocea* revealed 12,815 large, shared syntenic blocks, which encompassed 4,416 genes and 24 chromosomes (Table S7). The rearrangement events were distributed across all *A. latus* linkage groups, without evidence for local clustering (Fig. 2B). The most prominent event was the chromosomal fusion in *A. latus* (AL6 and AL20) that joined *L. crocea* chromosome LC2, thus revealing that some of the genes were located within the breakpoints of reshuffling events. These probably occurred in the last common ancestor of *A. latus* and *L. crocea* approximately 119.92 million years ago (Fig. 3A).

**Figure 3.**
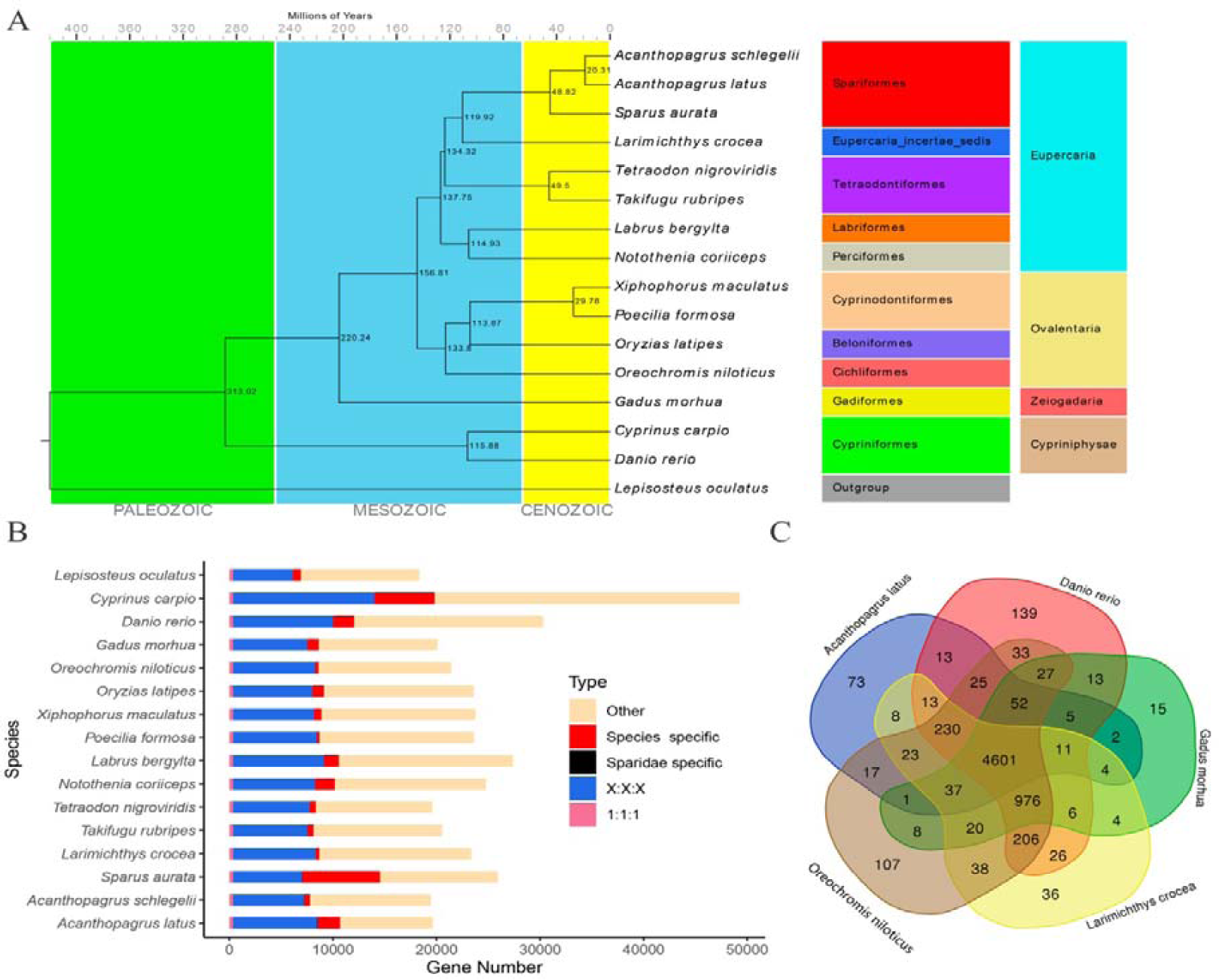
Comparative genomics analysis of *A. latus* and other teleosts. (A) Phylogenetic placement of *A. latus* in the teleosts phylogenetic tree which used 367 single-copy orthologous genes. The estimated divergence times are displayed below the phylogenetic tree. (B) Comparison of the gene repertoire of 16 teleosts genomes. “1:1:1” indicates universal single-copy genes; “X:X:X” indicates orthologues exist in all representative genomes (missing in one species not allowed), with “X” meaning one or more orthologs per species; “Sparidae specific” indicates genes only exist in the Sparidae fish. “Teleost specific” indicates orthologues exist in all teleost genomes. “Others” indicates orthologues genes that do not fit into the other categories. (C) The shared and unique gene families in five teleost fish are shown in the Venn diagram.

### 3.4 Phylogenetic relationships between *A. latus* and other vertebrates

To investigate the phylogenetic relationships of *A. latus* with other species, the genome of *A. latus* and the published genomes of 15 other vertebrate species were compared. We identified a total of 367 single-copy orthologous genes. Using those single-copy orthologues, we constructed a phylogenetic tree with 15 other fish species including *P. major, A. schlegelii, S. aurata, L. crocea, L. bergylta, N. coriiceps, T. rubripes, T. nigroviridis, X. maculatus, P. formosa, O. latipes, O. niloticus, G. morhua, C. carpio, D. rerio*, and *L. oculatus. L. oculatus* was used as an outgroup and served as a basis for investigating the evolutionary trajectory of *A. latus* (Fig. 3A). *A. latus* was placed in family Sparidae, which was consistent with the above sequencing read blast results. We found that *A. latus* diverged ∼72.53 million years ago from other Sparidae fish (*P. major, S. cantharus*, and *S. aurata*) and ∼155.69 million years ago from its common ancestor with *L. crocea* (Fig. 3A).

Compared with other teleost genomes, the *A. latus* genome has 35 Sparidae-specific genes and 2,170 species-specific genes (>11.0% of the entire gene repertoire, Fig. 3B, Table S8), including genes related to cellular processes, environmental information processing, genetic information processing, metabolism and organismal systems (Fig. S3, Table S9).

Moreover, a five-way orthologue comparison among related teleosts (*A. latus, D. rerio, G. morhua, L. crocea*, and *O. niloticus*) revealed that 4,601 gene families were shared by all five species (Fig. 3C), accounting for 89.95% of all *A. latus* gene families. A total of 73 were specific to *A. latus*.

### 3.5 Gene family comparisons

Gene family expansion is arguably one of the most important contributors to phenotypic diversity and evolutionary adaptions to the environment (Rayna & Hans, 2015). We compared the gene families of *A. latus* and eight other teleost species including *S. aurata, D. rerio, G. morhua, L. oculatus, O. niloticus, O. latipes, P. formosa*, and *T. rubripes* and identified 238 expanded (*p* < 0.05) and 87 contracted (*p* < 0.05) gene families (Table S10) with fewer than 100 gene copies in *A. latus* (Fig. 4A). Moreover, only 1 expanded (*p* > 0.05) gene family and 5 contracted (*p* > 0.05) gene families (Table S11) were detected with more than 100 gene copies in *A. latus*. In addition, the GO annotations of the expanded genes were indicative of their involvement in integral component of membrane (GO: 0016021), zinc ion binding (GO: 0008270), metal ion binding (GO: 0046872), nucleic acid binding (GO: 0003676), and intracellular (GO: 0005622) functions were found (Fig. S4) and their significant enrichment were shown in 34 GO terms and 104 KEGG pathways (Fig. S5). In particular, the expanded gene families were overrepresented in the NOD-like receptor signalling pathway (map04621), cellular senescence pathway (map04218), and cell adhesion molecules (map04514).

**Figure 4.**
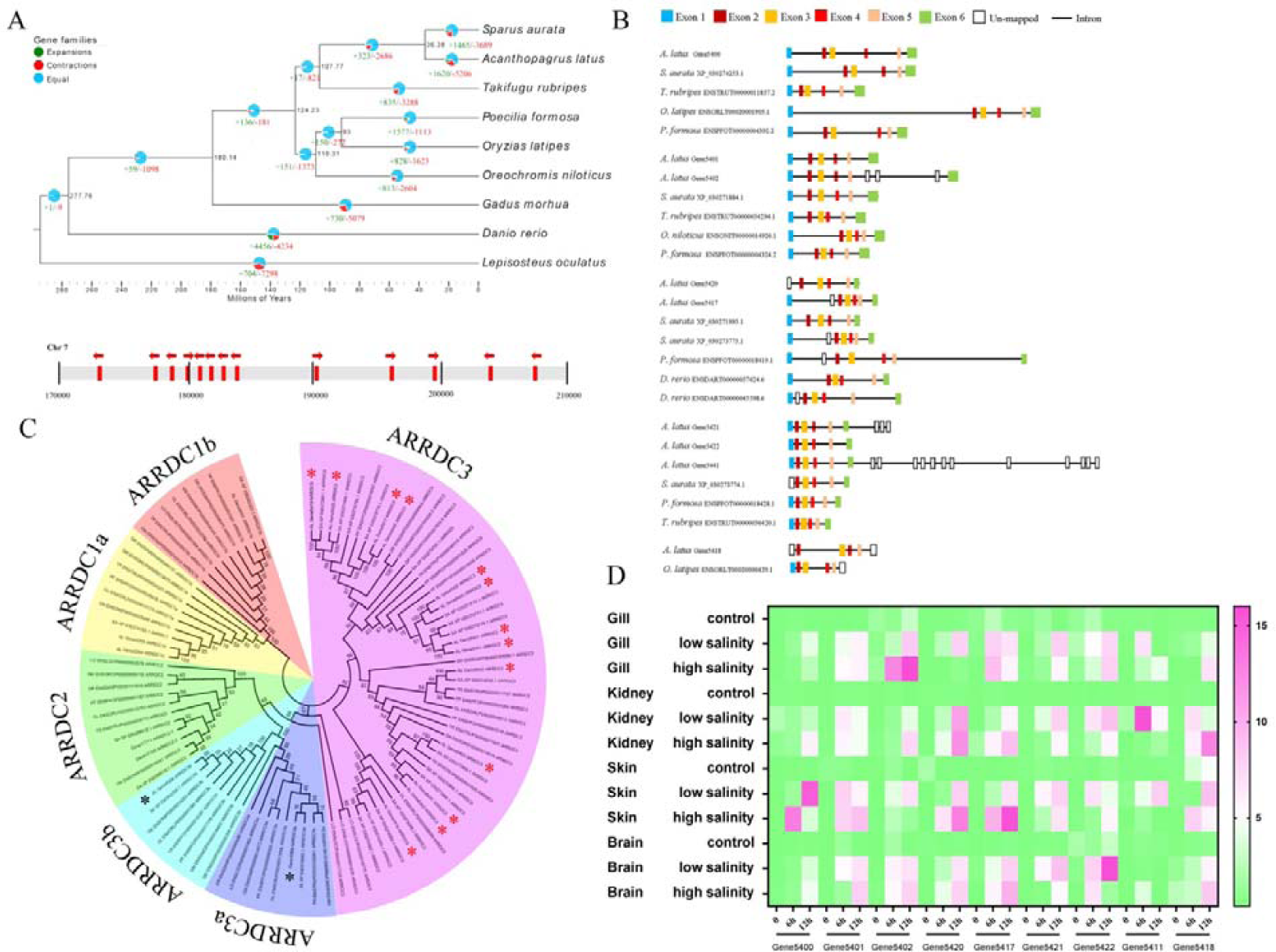
ARRDC gene family in *A. latus*. (A) Dynamic evolution of gene families among nine teleost species. Green and red numbers represent the expanded or contracted gene families in each linage, respectively. MRCA: most recent common ancestor. (B) Phylogenetic tree of the ARRDC gene family in teleosts. Six clades of ARRDC in the *A. latus* genome were observed, the expanded genes are marked by red stars. The genes in largest clade are specifically expanded *ARRDC3* in the *A. latus* genome, which are also tandemly duplicated. The arrow indicates the transcriptional orientation. (C) Structure of the *ARRDC3* gene in teleosts. (D) Temporal expression of nine *ARRDC3* genes in gill, kidney, skin, and brain after acute salinity stress (low and high salinity) for the indicated time points.

For these important pathways, we found extreme expansion of the arrestin domain containing protein 3 (ARRDC3), glutathione-S-transferase A (GSTA), and solute carrier family 12 member (SLC12A) in the genomes of two euryhaline Sparidae species (*A. latus* and *S. aurata*), while very few were detected in the other seven teleost fish. ARRDCs are important for regulating signal transduction within cells (Huo et al., 2015). Mice and humans have at least five a-arrestins: ARRDC1-4 and thioredoxin-interacting protein (TXNIP). Previous studies have revealed that a-arrestins are a family of proteins with roles in regulating metabolism and the development of obesity (Patwari et al., 2011; Ogawa et al., 2019). However, the function of ARRDCs has not been reported for fish. We speculate that ARRDCs may play a role in osmoregulation and metabolism, especially the expanded genes. In the present study, to identify ADDRC3 in the osmoregulation signalling pathway and to study its evolutionary history, we used a comparative and phylogenetic approach involving a large number of vertebrate species. A total of 94 ADDRC proteins from 9 fish were used to construct a phylogenetic tree, and three major clades were resolved, corresponding to *ARRDC1, ARRDC2*, and *ARRDC3*, which included ARRDC3a, ARRDC3b, and ARRDC3 (Fig. 4B). Two Sparidae fish, *A. latus* and *S. aurata*, possessed 13 and 16 *ARRDC3* genes, respectively, which were greater than the numbers in other species. These genes were determined to be tandemly duplicated, and these duplicates shared high sequence similarities (identity >98%), thus indicating recent duplications. However, *ARRDC3* was not identified in cod, a marine benthic fish. Additionally, we found that *A. latus* had two *ARRDC1a* paralogous genes; however, there was only one in other species.

Generally, structural complexity can be caused by exon gains or losses, which is a core evolutionary mechanism in most gene families (Yu et al., 2018). The genomic structure of *ARRDC3* was divided into five types based on comparisons of amino acid coding sequences among the 9 teleosts (Fig. 4C). The diversity of the *ARRDC3* genomic structure is caused by increased coding sequences, suggesting that exon gain was the evolutionary mechanism of *ARRDC3* in the two euryhaline species. Furthermore, the results of the acute salinity stress experiment showed that the expansion of *ADDRC3* genes responds rapidly to salinity changes, varying significantly within 6 h or 12 h subsequent to low or high salinity stress in the osmoregulatory tissues (gill, kidney, skin, and brain) (Fig. 4D), suggesting that the *ADDRC3* gene family is involved in the adaptation mechanism of salinity conversion.

Conversion in response to salinity can cause an increase in reactive oxygen species (ROS) in fish that may result in oxidative damage if not cleared quickly (Martínez-Álvarez et al., 2005). Antioxidant mechanisms are the main barrier against oxidative damage in fish. Fish can scavenge ROS and enhance resistance through antioxidant mechanisms (Rudneva 1997). GST is an important enzyme in the antioxidant defence system and can remove free radicals and active substances by catalysis (Güven et al., 2010; Gonçalves-Soares et al., 2012). Salinity stress can also induce GST activity in the liver of *O. niloticus* and in the gill of *Paralichthys olivaceus* (Japanese flounder) (Choi et al., 2008; El-Leithy et al., 2019). In the present study, a total of 54 GST proteins from 9 fish were used to construct a phylogenetic tree, and three major clades were resolved, corresponding to *GSTA, GSTT*, and *GSTZ* (Fig. S6). We found prominent expansion of *GSTA* genes in the genomes of the euryhaline fish *A. latus* and *S. aurata*, while very few paralogous genes were observed in other teleosts. A typical genome structure of *GSTA* was displayed in Fig. S7. It showed that the loss of one or two exons was the evolutionary mechanism of *GSTA* in *A. latus* (Yu et al., 2018).

Expansion of *GSTA* genes was also observed in the genomes of the two euryhaline fishes, yellowfin seabream and gilthead seabream. Additionally, the expanded genes of *GSTA* responded rapidly to salinity changes and varied significantly within 6 h or 12 h subsequent to low or high salinity stress in gill, kidney, skin, and brain (Fig. S8). These data provide strong support for the correlation of differential expression between the types of antioxidant defence genes with the habits and characteristics of euryhalinity, suggesting that those two seabream have a highly effective osmoregulatory control system that is adapted to their habitat transitions.

Additionally, the Na^+^/K^+^/2Cl^−^ cotransporter (NKCC), an SLC12A-family protein that transports Na^+^, K^+^ and 2Cl^−^ into cells, is essential for cell ionic and osmotic regulation (Xu et al., 2017). NKCCs play a crucial role in the homeostasis of cell volume and maintenance of electrolyte content and are involved in regulating transepithelial ion and water movement in polarized cells (Markadieu and Delpire, 2014). The SLC12A5- and SLC12A2 (NKCC1)-family genes, which are Na^+^/K^+^/2Cl^−^ cotransporters, are also significantly expanded in the yellowfin seabream genome (Fig. S9), and most of them are highly expressed in the gill, kidney, and skin after acute salinity changes (Fig. S10). These features might enable the seabream to rapidly perform signal transduction, thus enhancing its responsiveness and strengthening its osmoregulation, including energy metabolism, antioxidant defence, and ion transport.

### 3.6 Gonad development

Sexual dimorphism is widespread in nature and describes the complete set of differences between sexes (Jazin and Cahill, 2010; Gamble and Zarkower, 2012). Despite differences in morphology and behaviour, females and males are nearly genetically identical. However, many genes are more actively transcribed in one sex than in the other, and sex-biased expression is considered to be the manner by which sexual dimorphism can arise from the genome (Ranz et al., 2003; Ellegren and Parsch, 2007; Parsch and Ellegren, 2013). Moreover, teleost fishes show great variations in how sex phenotypes form. Among them, Sparidae, which might be considered a model family, display a remarkable diversity of reproductive modes. In the present study, to explain the genetic basis of the gonads, their formation, and their function in *A. latus*, we performed transcriptome analyses of gonads among three periods (stage 1: male, stage 2: intersexual, and stage 3: female) (Fig. 5A). Compared with stage 1, 3,442 and 822 differentially expressed genes (DEGs) were upregulated in stage 2 and stage 3, respectively. Additionally, 743 genes were significantly upregulated in both stage 2 and stage 3. KEGG analyses indicated that many of these genes are involved in metabolic pathways (map01100), apoptosis (map04210), biosynthesis of secondary metabolites (map01110), and tight junctions (map04530) (Fig. S11). Specifically, 17 genes encode proteins related to gonad development, containing the cytochrome p450 family, forkhead box protein, and sox family (Fig. 5B). However, those genes play important roles in the regulation of sex reversal, as indicated in previous studies (Wei et al., 2016; Cao and Chen, 2019; Fan et al., 2019b). We speculate that the regulation of sex reversal in *A. latus* is related to those 17 genes.

**Figure 5.**
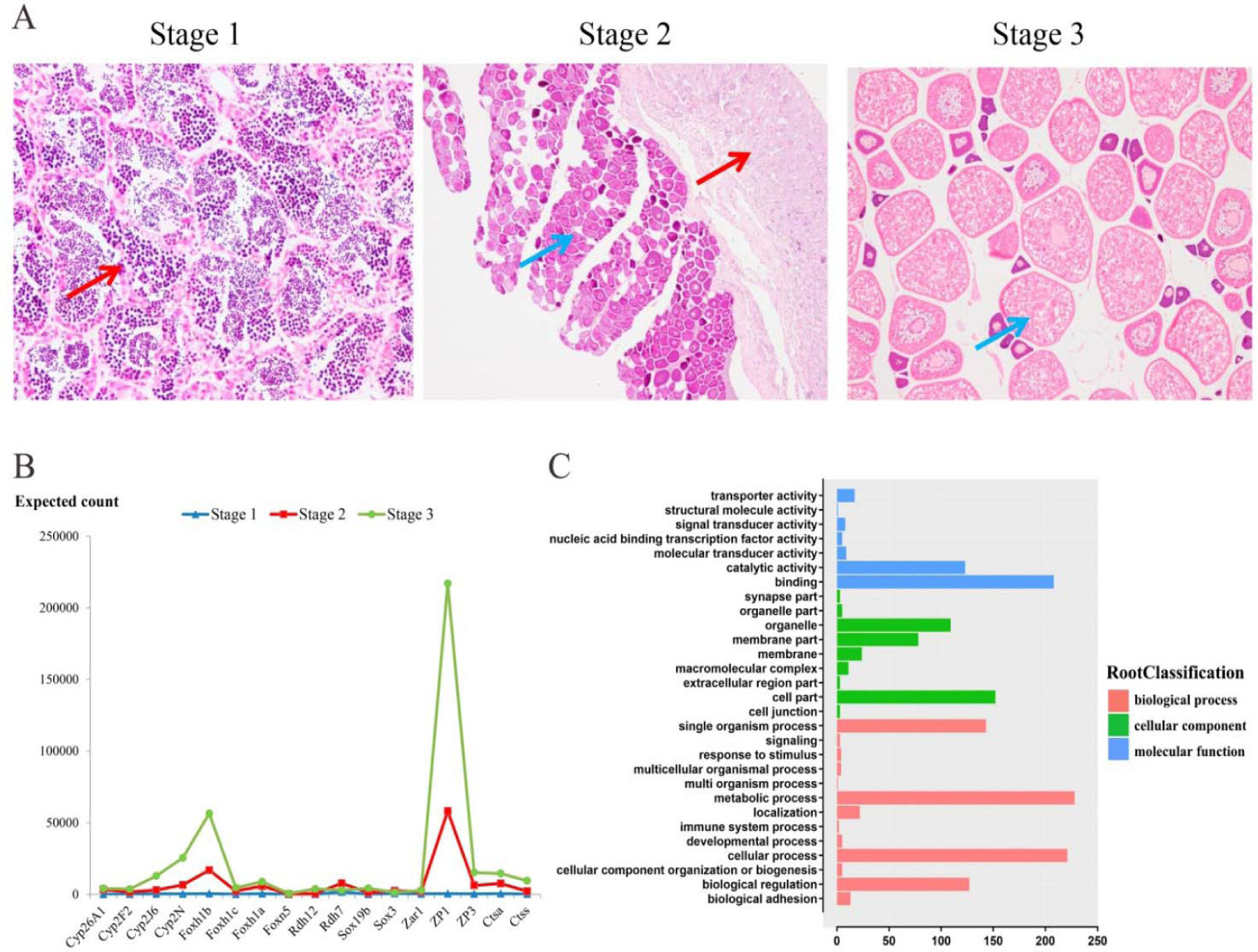
Regulation of genes related to sex reversal and function identified from developmental stages and adult tissues transcriptome data. (A) Different stages of gonad development in adult tissues. Stage 1 indicates male gonad, stage 2 indicates intersexuality gonad, stage 3 indicates female gonad. Red and blue arrows indicate male and female gonad tissues, respectively. (B) The gene expression pattern involved in sex reversal–regulated genes in gonad developmental stages. (C) Distribution of gonad development-specific genes in GO annotations indicative of abundance in binding, metabolic process, and cellular process.

To further confirm the numbers of specific genes in the three stages, compared with stage 2 and stage 3, 184 genes were specifically expressed in stage 1. Compared with stage 1 and stage 3, 146 genes were specifically expressed in stage 2. Moreover, compared with stage 1 and stage 2, 182 genes were specifically expressed in stage 3. In addition, according to GO annotations, 2,261 gonad development-specific genes were associated with binding, metabolic process, cellular process, cell part, biological regulation, and signal organism process functions (Fig. 5C). To further evaluate the reliability of RNA sequencing, 30 differentially expressed genes and 30 differentially expressed mRNAs (10 genes in each stage) were randomly selected to validate their relative expression levels in gonad development using qRT-PCR (Fig. S12). As shown in Fig. S12, the trend of the differential expression levels of these 30 randomly selected genes was 100% consistent with the pattern in our RNA-seq data, indicating that the RNA-seq data are reliable.

Transcription factors such as TGF-β1, TGF-β3, Smad, growth differentiation factor 9, Amh/amh, and follistatin may be important for gonadal differentiation in fish (Memon et al., 2008; Garrel et al., 2016; Chen et al., 2017). Several genes (TGF beta, TGF beta precursor, BMP1-8, Amh, Smad1-8, Follistatin, transforming protein RhoA-like, and transcription factor E2F4-like) are differentially expressed in the transforming growth factor beta (TGF-β) signalling pathway, thus suggesting that these genes may be involved in sex differentiation and development in *A. latus*. We also find that the Wnt signalling pathway, which plays a pivotal role in signal transduction and cell proliferation, contained sex differentiation- and development-related genes (Frizzled, Wnt1-11, RhoA, DVL) (Stanimirovic et al., 2018; Luo et al., 2019; Yuan et al., 2019).

In summary, we have completed a chromosome genome assembly of *A. latus*. The continuity of both the contig and scaffold sequences, represented by their N50 lengths, reached high levels comparable to those in other model teleosts. More than 92% of BUSCO genes were identified in the genome, which implies the completeness of the genome. In total, 19,631 protein-coding genes were predicted and functionally annotated in the *A. latus* genome. The phylogenetic relationships between *A. latus* and other closely related teleosts were analysed. This study provides the first chromosome-level genome of the Sparidae. This high-quality genome provides a solid foundation for future population and conservation studies of Sparidae fish, as well as for investigations of environmental adaptation in fish native to euryhaline environments and with sex reversal.

## ACKNOWLEDGEMENTS

This work was supported by Financial Fund of Ministry of Agriculture and Rural affairs of China (NHYYSWZZZYKZX2020), Key Special Project for Introduced Talents Team of Southern Marine Science and Engineering Guangdong Laboratory (Guangzhou) (GML2019ZD0605), Natural Science Foundation of Guangdong Province (2017A030310594), China-ASEAN maritime cooperation fund (00-201620821), National Infrastructure of Fishery Germplasm Resources Project (2019DKA30470) and Science and Technology Infrastructure Construction Project of Guangdong Province (2019B030316030).

## AUTHOR CONTRIBUTIONS

K.C.Z., S.G.J., and D.C.Z. designed the research and wrote the paper. K.C.Z. performed the research. H.Y.G. and N.Z. analyzed the data. B.S.L. and L.G. contributed reagents/materials/analysis tools.

## CONFLICTS OF INTEREST

The authors declare no competing financial interests.

## DATA AVAILABILITY

The raw genome and RNA sequencing data have been deposited in the SRA under Bioproject number PRJNA566024. This Whole Genome Shotgun project has been deposited at DDBJ/ENA/GenBank under the accession JABAYC000000000.

## Notes

### Competing Interest Statement

The authors have declared no competing interest.

## REFERENCES

Bao, L.S., Tian, C.X., Liu, S.K., Zhang, Yu., Elaswad, A., et al. (2019). The Y chromosome sequence of the channel catfish suggests novel sex determination mechanisms in teleost fish. BMC Biology, 17, 6. https://doi.org/10.1186/s12915-019-0627-7

Bao, W., Kojima, K. K., & Kohany, O. (2015). Repbase Update, a database of repetitive elements in eukaryotic genomes. Mobile DNA, 6, 11. https://doi.org/10.1186/s13100-015-0041-9.

Bauchot, M. L., & Smith, M. M. (1986). Sparidae. In FAO Species Identification Sheets for Fisheries Purposes Western Indian Ocean. Fishing Area, 51, (4), 1–11.

Benson, G. (1999). Tandem repeats fnder: a program to analyze DNA sequences. Nucleic Acids Research, 27, 573–580. https://doi.org/10.1093/nar/27.2.573

Birney, E., Clamp, M., & Durbin, R. (2004). GeneWise and Genomewise. Genome Research, 14, 988–995. https://doi.org/10.1101/gr.1865504

Boeckmann, B., Bairoch, A., & Apweiler, R. (2003). The SWISS-PROT protein knowledgebase and its supplement TrEMBL in 2003. Nucleic Acids Research, 31(1), 365–370. https://doi.org/10.1093/nar/gkg095

Burton, J. N., Adey, A., Patwardhan, R. P., Qiu, R., Kitzman, J. O., & Shendure, J. (2013). Chromosome-scale scaffolding of de novo genome assemblies based on chromatin interactions. Nature Biotechnology, 31(12), 1119–1125. https://doi.org/10.1038/nbt.2727

Buxton, C. D., & Garratt, P. A. (1990). Alternative reproductive styles in seabreams (Pisces: Sparidae). Environmental Biology of Fishes, 28, 113–124. https://doi.org/10.1007/bf00751031

Cao, J., & Cheng, X. Z. (2019). Transcriptome-based identification and molecular evolution of the cytochrome p450 genes and expression profiling under dimethoate treatment in Amur Stickleback (*Pungitius sinensis*). Animals, 9(11), 873. https://doi.org/10.3390/ani9110873

Carpenter, K. E. (2001). Family Sparidae. In Species Identification Guide for Fishery Purposes. The living marine resources of the western central pacific, 5(3), 2990–3003.

Chen, S.L., Zhang, G.J., Shao, C.W., Huang, Q.F., Liu, G., Zhang, P., Song, W.T., An, N., Chalopin, D., Volff, J.N., et al. (2014). Whole-genome sequence of a flatfish provides insights into ZW sex chromosome evolution and adaptation to a benthic lifestyle. Nature genetics, 46(3), 253. https://doi.org/10.1038/ng.2890

Chen, W., Liu, L., & Ge, W. (2017). Expression analysis of growth differentiation factor 9 (Gdf9/gdf9), anti-müllerian hormone (Amh/amh) and aromatase (Cyp19a1a/cyp19a1a) during gonadal differentiation of the zebrafish, *Danio rerio*. Biology of Reproduction, 96(2), 401–413. https://doi.org/10.1095/biolreprod.116.144964

Chen, Y., Hong, W. S., Wang, Q., & Chen, S. X. (2015). Cloning and expression pattern of gsdf during the first maleness reproductive phase in the protandrous *Acanthopagrus latus*. General and Comparative Endocrinology, 217-218, 71-80. https://doi.org/10.1016/j.ygcen.2015.02.018

Choi, C. Y., An, K. W., & An, MI. (2008). Molecular characterization and mRNA expression of glutathione peroxidase and glutathione S-transferase during osmotic stress in olive flounder (*Paralichthys olivaceus*). Comparative Biochemistry and Physiology A-molecular & integrative physiology, 149(3), 330–337. https://doi.org/10.1016/j.cbpa.2008.01.013

Cook, D. G., & Herbert, N. A. (2012). Low O-2 avoidance is associated with physiological perturbation but not exhaustion in the snapper (*Pagrus auratus*: Sparidae). Comparative Biochemistry and Physiology A-molecular & integrative physiology, 162(4), 310–316. https://doi.org/10.1242/jeb.073023

Cox, M. P., Peterson, D. A., & Biggs, P. J. (2010). SolexaQA: At-a-glance quality assessment of Illumina second-generation sequencing data. BMC bioinformatics, 11, 485. https://doi.org/10.1186/1471-2105-11-485

Chen, X., Jiang, Y., Gao, F., Zheng, W., Krock, T.J., Stover, N.A., Lu, C., Katz, L.A., Song, W. (2019). Genome analyses of the new model protist euplotes vannus focusing on genome rearrangement and resistance to environmental stressors. Molecular Ecology Resources, 19 (5), 1292–1308. https://doi.org/10.1111/1755-0998.13023.

Cui, Z., Liu, Y., Wang, W., Wang, Q., Zhang, N., Lin, F., Wang, N., Shao, C., Dong, Z., Li, Y., et al. (2017). Genome editing reveals dmrt1 as an essential male sex-determining gene in Chinese tongue sole (*Cynoglossus semilaevis*). Scientific Reports, 7, 42213. https://doi.org/10.1038/srep42213

De Bie, T., Cristianini, N., Demuth, J. P., & Hahn, M. W. (2006). CAFE: A computational tool for the study of gene family evolution. Bioinformatics, 22(10), 1269–1271. https://doi.org/10.1093/bioinformatics/btl097

Dudchenko, O., Batra, S. S., Omer, A. D., Nyquist, S. K., Hoeger, M., … Durand, N.C. (2017). De novo assembly of the genome using Hi-C yields chromosome-length scaffolds. Science, 356, 92. https://doi.org/10.1126/science.aal3327

Edgar, R. C. (2004). MUSCLE: multiple sequence alignment with high accuracy and high throughput. Nucleic acids research, 32, 1792–1797. https://doi.org/10.1093/nar/gkh340

Ellegren, H., & Parsch, J. (2007). The evolution of sex-biased genes and sex-biased gene expression. Nature Reviews Genetics, 8, 689–698. https://doi.org/10.1038/nrg2167

El-Leithy, A. A. A., Hemeda, S. A., El-Naby, W. S. H. A., El-Nahas, A. F., Hassan S.A.H., Awad, S.T., El-Deeb. S.I., … Helmy, Z.A. (2019). Optimum salinity for Nile tilapia (Oreochromis niloticus) growth and mRNA transcripts of ion-regulation, inflammatory, stress-and immune-related genes. Fish Physiology and Biochemistry, 45(4), 1217–1232. https://doi.org/10.1007/s10695-019-00640-7

Fan, G.Y., Zhang, Y.L., Liu, X.C., Wang, J.H., Sun, Z.G., et al., (2019a). The first chromosome-level genome for a marine mammal as a resource to study ecology and evolution. Molecular Ecology Resources, 19 (4), 944–956. https://doi.org/10.1111/1755-0998.13003.

Fan, Z. F., Zou, Y. X., Liang, D. D., Tan, X. G., Jiao, S., Wu, Z.H., Zhang, P. J., … You, F. (2019b). Roles of forkhead box protein L2 (foxl2) during gonad differentiation and maintenance in a fish, the olive flounder (*Paralichthys olivaceus*). Reproduction Fertility and Development, 31(11), 1742–1752. https://doi.org/10.1071/RD18233

Gamble, T., & Zarkower, D. (2012). Sex determination. Current Biology, 22, 257–262. http://doi.org/10.1016/j.cub.2012.02.054

Garrel, G., Racine, C., L’Hote, D., Denoyelle, C., Guigon, C. J., di Clemente, N., & Cohen-Tannoudji, J. (2016). Anti-Mullerian hormone: a new actor of sexual dimorphism in pituitary gonadotrope activity before puberty. Scientific Reports, 6, 23790. https://doi.org/10.1038/srep23790

Gonçalves-Soares D., Zanette J., Yunes J. S., Yepiz-Plascencia G. M., & Bainy A. C. D. (2012). Expression and activity of glutathione S-transferases and catalase in the shrimp *Litopenaeus vannamei* inoculated with a toxic *Microcystis aeruginosa* strain. Marine Environmental Research, 75, 54–61. https://doi.org/10.1016/j.marenvres.2011.07.007

Guindon, S., & Gascuel, O. (2003). A simple, fast, and accurate algorithm to estimate large phylogenies by maximum likelihood. Molecular Systems Biology, 52, 696–704. https://doi.org/10.1080/10635150390235520

Guindon, S., Dufayard, J. F., & Lefort, V. (2010). New algorithms and methods to estimate maximum-likelihood phylogenies: assessing the performance of PhyML 3.0. Molecular Systems Biology, 59, 307–321. https://doi.org/10.1093/sysbio/syq010

Güven, A. (2010). The effect of kefir on the activities of GSH-Px, GST, CAT, GSH and LPO levels in carbon tetrachloride-induced mice tissues. Journal of Veterinary Medicine, 50(8), 412–416. https://doi.org/412-416.10.1046/j.1439-0450.2003.00693.x

Hattori, R.S., Murai, Y., Oura, M., Masuda, S., Majhi, S.K., Sakamoto, T., et al. (2012). A Y-linked anti-Mullerian hormone duplication takes over a critical role in sex determination. Proc Natl Acad Sci U.S.A. 109, 2955–2959. https://doi.org/10.1073/pnas.1018392109

He, Y., Chang, Y., Bao, L.S., Yu, M.J., Li, R., et al., (2019). A chromosome-level genome of black rockfish, *sebastes schlegelii*, provides insights into the evolution of live birth. Molecular Ecology Resources, 19 (5), 1309–1321. https://doi.org/10.1111/1755-0998.13034

He, Z., Zhang, H., Gao, S., Lercher, M. J., Chen, W. H., & Hu, S. (2016). Evolview v2: an online visualization and management tool for customized and annotated phylogenetic trees. Nucleic Acids Research, 44, 236–241. https://doi.org/10.1093/nar/gkw370

Hesp, S. A., Potter, I. C., & Hall, N. G. (2004). Reproductive biology and protandrous hermaphroditism in *Acanthopagrus latus*. Environmental Biology of Fishes, 70, 257–272. https://doi.org/10.1023/b:ebfi.0000033344.21383.00

Imakaev, M., Fudenberg, G., Mccord, R. P., Naumova, N., Goloborodko, A., & Lajoie, B. R. (2012). Iterative correction of Hi-C data reveals hallmarks of chromosome organization. Nature Methods, 9(10), 999. https://doi.org/10.1038/NMETH.2148

Jaffer, Y. D., Saraswathy, R., Ishfaq, M., Antony, J., Bundela, D. S., & Sharma, P. C. (2020). Effect of low salinity on the growth and survival of juvenile pacific white shrimp, Penaeus vannamei: A revival. Aquaculture, 515, 734561.

Jazin, E., & Cahill, L. (2010). Sex differences in molecular neuroscience: from fruit flies to humans. Nature Reviews Neuroscience, 11, 9–17. https://doi.org/10.1038/nrn2754

Kajitani, R., Toshimoto, K., Noguchi, H., Toyoda, A., Ogura, Y., & Okuno, M. (2014). Efficient de novo assembly of highly heterozygous genomes from whole-genome shotgun short reads. Genome Research, 24, 1384–1395. https://doi.org/10.1101/gr.170720.113

Kamiya, T., Kai, W., Tasumi, S., Oka, A., Matsunaga, T., Mizuno, N., Fujita, M., Suetake, H., Suzuki, S. & Hosoya, S., et al. (2012). A trans-species missense SNP in Amhr2 is associated with sex determination in the tiger pufferfish, *Takifugu rubripes* (Fugu). PLoS Genetics, 8(7), e1002798. https://doi.org/10.1371/journal.pgen.1002798

Kim, D., Langmead, B., & Salzberg, S.L. (2015). HISAT: a fast spliced aligner with low memory requirements. Nature methods, 12(4), 357–360. https://doi.org/10.1038/nmeth.3317

Kisten, Y, Strydom, NA, & Perissinotto, R. (2019). The effects of hypersalinity on the growth and skeletal anomalies of juvenile Cape stumpnose, *Rhabdosargus holubi* (Sparidae). Scientia Marina, 83(1), 61–68.

Kültz, D. (2012). The combinatorial nature of osmosensing in fishes. Physiology, 27(4), 259–275. https://doi.org/10.1152/physiol.00014.2012

Kültz, D. (2015). Physiological mechanisms used by fish to cope with salinity stress. Journal of Experimental Biology, 218(12):1907–1914. https://doi.org/10.1242/jeb.118695.

Laiz-Carrión, R., Guerreiro, P. M., Fuentes, J., Canario, A. V. M., Martín del Rio, M. P., & Mancera, J. M., (2005). Branchial osmoregulatory response to salinity in the gilthead sea bream, *Sparus aurata*. Journal of Experimental Zoology Part A-Ecological Genetics and Physiology, 303, 563–576. https://doi.org/10.1002/jez.a.183

Langmead, B., & Salzberg, S. L. (2012). Fast gapped-read alignment with Bowtie 2. Nature Methods, 9, 357–359. https://doi.org/10.1038/nmeth.1923

Li, H., & Durbin, R., (2009). Fast and accurate short read alignment with Burrows–Wheeler transform. Bioinformatics, 25(14), 1754–1760. https://doi.org/10.1093/bioinformatics/btp324

Li, J. E. & Ou, Y. J. (2000). Studies on the reproductive biology of the pond-cultured *Sparus latus* Houttuyn in the Coast of Shenzhen Bay. Journal of Zhejiang Ocean University (Natural Science), 19, 139-143. (In Chinese)

Liu, L. S., Yang, J. H., Lin, J. H., & Hong, M. X. (1991). Study on chromosome type of yellowfin bream. Chinese Journal of Zoology, 26(1), 14–16.

Livak, K. J., & Schmittgen, T. D., (2001). Analysis of relative gene expression data using real-time quantitative PCR and the 2^−Δ^^ΔCT^ method. Methods, 25, 402–408. https://doi.org/10.1006/meth.2001.1262

Luo, X. L., Liu, Y. T., Ma, S. J., Liu, L., Xie, R., & Wang, S. C. (2019). WDR34 Activates Wnt/Beta-Catenin Signaling in Hepatocellular Carcinoma. Digestive Diseases and Sciences, 64(9), 2591–2599. https://doi.org/10.1007/s10620-019-05583-w

Marcais, G., Delcher, A. L., Phillippy, A. M., Coston, R., Salzberg, S. L., & Zimin, A. (2018). MUMmer4: A fast and versatile genome alignment system. PLoS Computational Biology, 14, e1005944. https://doi.org/10.1371/journal.pcbi.1005944

Markadieu, N., & Delpire, E., (2014). Physiology and pathophysiology of SLC12A1/2 transporters. Pflugers Arch, 466, 91–105. https://doi.org/10.1007/s00424-013-1370-5

Marshall, W. S. (2013). Osmoregulation in estuarine and intertidal fishes. Fish physiology, 32, 395–434. https://doi.org/10.1016/B978-0-12-396951-4.00008-6

Martínez-Álvarez, RM., Morales, A. E., & Sanz, A. (2005). Antioxidant defenses in fish: biotic and abiotic factors. Reviews in Fish Biology & Fisheries, 15(1-2), 75-88. https://doi.org/10.1007/s11160-005-7846-4

Matsuda, M., Nagahama, Y., Shinomiya, A., Sato, T., Matsuda, C., Kobayashi, T., et al. (2002). DMY is a Y-specific DM-domain gene required for male development in the medaka fish. Nature, 417, 559–563. https://doi.org/10.1038/nature751

Maziade, M., Bouchard, S., & Gingras, N. (1996). Long-term stability of diagnosis and symptom dimensions in a systematic sample of patients with onset of schizophrenia in childhood and early adolescence. II: Positive/negative distinction and childhood predictors of adult outcome. British Journal of Psychiatry, 169(03), 371–378.

Moriya, Y., Itoh, M., Okuda, S., Yoshizawa, A.C., & Kanehisa, M. (2007). KAAS: an automatic genome annotation and pathway reconstruction server. Nucleic Acids Research, 35, 182–185. https://doi.org/10.1093/nar/gkm321

Myosho, T., Otake, H., Masuyama, H., Matsuda, M., Kuroki, Y., Fujiyama, A., et al. (2012). Tracing the emergence of a novel sex-determining gene in Medaka, *Oryzias luzonensis*. Genetics 191, 163–170. https://doi.org/10.1534/genetics.111.137497

Nelson, J. S. (2006). Fishes of the World (Fourth Edition). New Jersey: John Wiley & Sons Inc, 371. https://doi.org/10.1111/j.1467-2979.2006.00227.x

Palaiokostas, C., Bekaert, M., Taggart, J., Gharbi, K., McAndrew, B., Chatain, B., et al. (2015). A new SNP-based vision of the genetics of sex determination in European sea bass (*Dicentrarchus labrax*). Genet. Sel. Evol. 47:68. https://doi.org/10.1186/s12711-015-0148-y.

Parsch, J., & Ellegren, H. (2013). The evolutionary causes and consequences of sex-biased gene expression. Nature Reviews Genetics, 14, 83–87. https://doi.org/10.1038/nrg3376

Pauletto, M.P., Tereza, M., Serena, F., Massimiliano, B., Alexandros, T., et al. (2018). Genomic analysis of Sparus aurata reveals the evolutionary dynamics of sex-biased genes in a sequential hermaphrodite fish. Communications Biology, 1, 119. https://doi.org/10.1038/s42003-018-0122-7

Ralf, J., Emil, A., Betnér E., Sifra, B., Will, F., & Lars, H. (2018). Embryonic expression patterns and phylogenetic analysis of panarthropod sox genes: insight into nervous system development, segmentation and gonadogenesis. BMC Evolutionary Biology, 18(1), 88. https://doi.org/10.1186/s12862-018-1196-z

Ranz, J. M., Castillo-Davis, C. I., Meiklejohn, C. D., & Hartl, D. L. (2003). Sexdependent gene expression and evolution of the Drosophila transcriptome. Science, 300, 1742–1745. https://doi.org/10.1126/science.1085881

Rao, S. S. P., Huntley, M. H., Durand, N. C., Stamenova, E. K., Bochkov, I. D., … Robinson, J. T. (2014). A 3D map of the human genome at kilobase resolution reveals principles of chromatin looping. Cell, 159, 1665–1680. https://doi.org/10.1016/j.cell.2014.11.021

Rudneva II. (1997). Blood antioxidant system of black sea elasmobranch and teleost. Comparative Biochemistry and Physiology Part C: Pharmacology, Toxicology and Endocrinology, 118(2), 255–260. https://doi.org/10.1016/S0742-8413(97)00111-4

Ruiz-Jarabo, I., Martos-Sitcha, J. A., Barragan-Mendez, C., Martinez-Rodriguez, G., Mancera, J. M., & Arjona, F. J. (2017). Gene expression of thyrotropin- and corticotrophin-releasing hormones is regulated by environmental salinity in the euryhaline teleost *Sparus aurata*. Fish Physiology and Biochemistry, 44(2), 615–628. https://doi.org/10.1007/s10695-017-0457-x

Servant, N., Varoquaux, N., Lajoie, B.R., Viara, E., Chen, C.J., et al. (2015). HiC-Pro: An optimized and flexible pipeline for Hi-C data processing. Genome Biology, 16(1), 259. https://doi.org/10.1186/s13059-015-0831-x

Shao, C., Li, C., Wang, N., Qin, Y., Xu, W., & Liu, Q. (2018). Chromosome-level genome assembly of the spotted sea bass, *Lateolabrax maculatus*. GigaScience, 7(11). https://doi.org/10.1093/gigascience/giy114.

Shao, C.W., Chen, S.L., Scheuring, C.F., Xu, J.Y., Sha, Z.X., Dong, X.L., et al. (2010). Construction of Two BAC libraries from halfsmooth tongue sole *cynoglossus semilaevis* and identification of clones containing candidate sex-determination genes. Marine Biotechnology 12, 558–568. https://doi.org/10.1007/s10126-009-9242-x.

Simao, F. A., Waterhouse, R. M., Ioannidis, P., Kriventseva, E. V., & Zdobnov, E. M. (2015). BUSCO: Assessing genome assembly and annotation completeness with single-copy orthologs. Bioinformatics, 31(19), 3210–3212. https://doi.org/10.1093/bioinformatics/btv351

Stanimirovic, J., Obradovic, M., Panic, A., Petrovic, V., Alavantic, D., Melih, I., & Isenovic, E. R., (2018). Regulation of hepatic Na^+^/K^+^-ATPase in obese female and male rats: involvement of ERK1/2, AMPK, and Rho/ROCK. Molecular and Cellular Biochemistry, 440(1-2), 77–88. https://doi.org/10.1007/s11010-017-3157-z

Tamura, K., Stecher, G., & Peterson, D., (2013). MEGA 6: Molecular evolutionary genetics analysis version 6.0. Molecular Biology and Evolution, 3, 2725–2729. https://doi.org/10.1186/s12985-019-1247-0

Tarailo-Graovac, M., & Chen, N. (2009). Using RepeatMasker to identify repetitive elements in Genomic sequences. Current Protocols in Bioinformatics, 4, 4–10. https://doi.org/10.1002/0471250953.bi0410s25

Tinacci, L., Giusti, A., Guardone, L., Luisi, E., & Armani, A. (2019). The new Italian official list of seafood trade names (annex I of ministerial decree n. 19105 of September the 22nd, 2017): Strengths and weaknesses in the framework of the current complex seafood scenario. Food Control, 96, 68–75. https://doi.org/10.1016/j.foodcont.2018.09.002

Tsakogiannis, A., Manousaki, T., Lagnel, J., Papanikolaou, N., Papandroulakis, N., Mylonas, C. C., & Tsigenopoulos, C. S. (2019). The Gene Toolkit Implicated in Functional Sex in Sparidae Hermaphrodites: Inferences From Comparative Transcriptomics. Frontiers in Genetics, 9, 749. https://doi.org/10.3389/fgene.2018.00749

Wei, L., Yang, C., Tao, W.J., Wang, D.S. (2016). Genome-wide identification and transcriptome-based expression profiling of the sox gene family in the nile tilapia (*Oreochromis niloticus*). International Journal of Molecular Sciences, 17 (3), 270. https://doi.org/10.3390/ijms17030270.

Xu, B. P., Tu, D. D., Yan, M. C., Shu, M. A., & Shao, Q. J. (2017). Molecular characterization of a cDNA encoding Na^+^/K^+^/2Cl(^-^) cotransporter in the gill of mud crab (*Scylla paramamosain*) during the molt cycle: Implication of its function in osmoregulation. Comparative biochemistry and physiology a-molecular & integrative physiology, 203, 115–125. https://doi.org/10.1016/j.cbpa.2016.08.019

Xu, Z. X., Gan, L., Li. T. Y., Xu, C., Chen, K., Wang, X. D., Qin, J. G., Chen, L. Q., & Li, E. C. (2015). Transcriptome profiling and molecular pathway analysis of genes in association with salinity adaptation in Nile tilapia, *Oreochromis niloticus*. PLoS One 10(8), e0136506. https://doi.org/10.1371/journal.pone.0136506

Yang, Z. (2007). PAML 4: phylogenetic analysis by maximum likelihood. Molecular Biology and Evolution, 24, 1586–1591.

Yang, Z., & Rannala, B. (2006). Bayesian estimation of species divergence times under a molecular clock using multiple fossil calibrations with soft bounds. Molecular Biology and Evolution, 223, 212–226. https://doi.org/10.1093/molbev/msj024

Yano, A., Guyomard, R., Nicol, B., Jouanno, E., Quillet, E., Klopp, C., et al. (2012). An immune-related gene evolved into the master sex-determining gene in rainbow trout, *Oncorhynchus mykiss*. Curr. Biol. 22, 1423–1428. https://doi.org/10.1016/j.cub.2012.05.045

Yuan, J. B., Gao, Y., Sun, L. N., Jin, S. J., Zhang, X. J., Liu, C. Z., Li, F. H., Xiang, & J. H., (2019). Wnt signaling pathway linked to intestinal regeneration via evolutionary patterns and gene expression in the sea cucumber *apostichopus japonicas*. Frontiers in Genetics, 10, 112. https://doi.org/10.1016/0014-4894(77)90131-x

Zhang, Z. Y., Zhang, K., Chen, S. Y., Zhang, Z. W., Zhang, J. Y., You, X. X., Bian, C., Xu, J., Jia, C. F., & Qiang, J. (2018). Draft genome of the protandrous Chinese black porgy, *Acanthopagrus schlegelii*. GigaScience, 7(4), 1. https://doi.org/10.1093/gigascience/giy012

Zhou, Q., Gao, H.Y., Zhang, Y., Fan, G.Y., Xu, H., Zhai, J.M., Xu, W.T., Chen, Z.F., Zhang, H., Liu, S.S., Niu, Y.P., Li, W.S., Li, W.M., Lin, H.R., Chen, S.L. (2019). A chromosome-level genome assembly of the giant grouper (*Epinephelus lanceolatus*) provides insights into its innate immunity and rapid growth. Molecular Ecology Resources, 19 (5), 1322–1332. https://doi.org/10.1111/1755-0998.13048.

Zhu, K. C., Liu, B. S., Guo, H. Y., Zhang, N., Guo, L., Jiang, S. G. & Zhang, D. C. (2020). Functional analysis of two MyoDs revealed their role in the activation of myomixer expression in yellowfin seabream (*Acanthopagrus latus*) (Hottuyn, 1782). International Journal of Biological Macromolecules, https://doi.org/10.1016/j.ijbiomac.2019.11.139

