## supplemental files for "A chromosome-level genome assembly of the yellowfin seabream (*Acanthopagrus latus*) (Hottuyn, 1782) provides insights into its osmoregulation and sex reversal"

**Supplementary Figures**

**Figure S1. The estimated genome size of female individual of *A. latus* is used by k-mer depth and frequency distribution.** Values for K-mers are plotted against the frequency (y axis) of their depth (x axis).

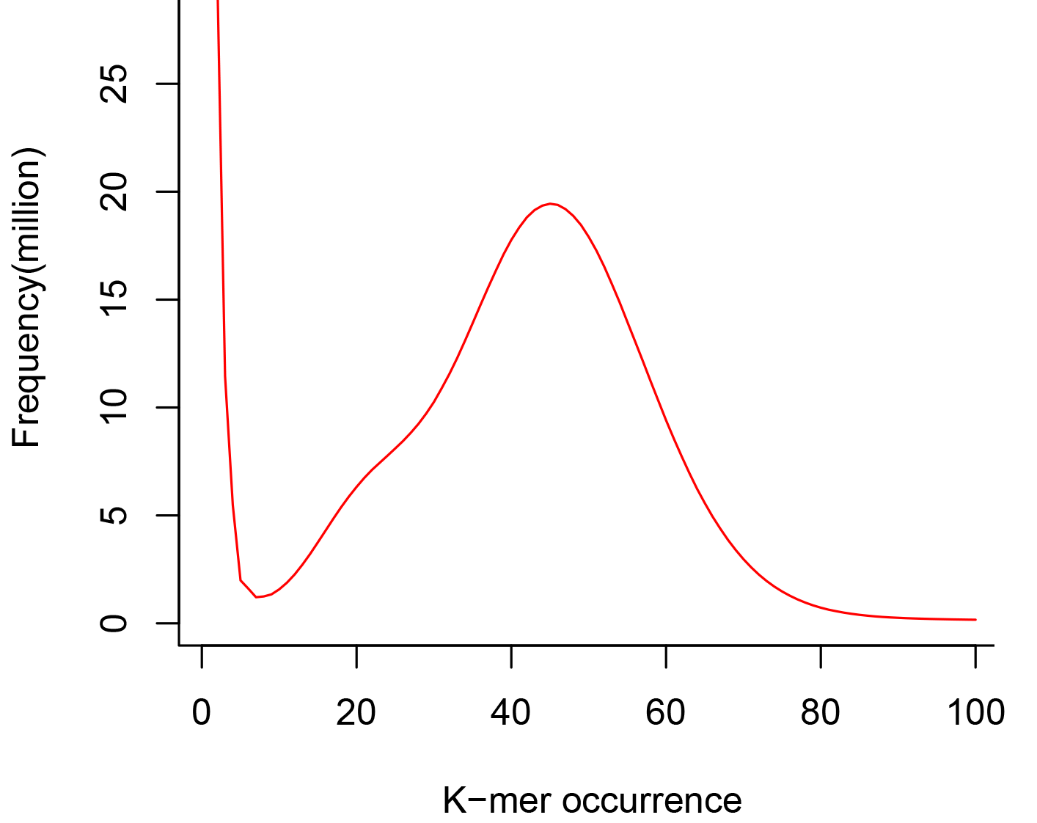

**Figure S2. Distribution of Insert size in six libraries.** (a) Insert distribution of 180 bp library; (b) Insert distribution of 500 bp library; (c) Insert distribution of 3 kb library; (d) Insert distribution of 5 kb library; (e) Insert distribution of 10 kb library; (f) Insert distribution of 14 kb library.

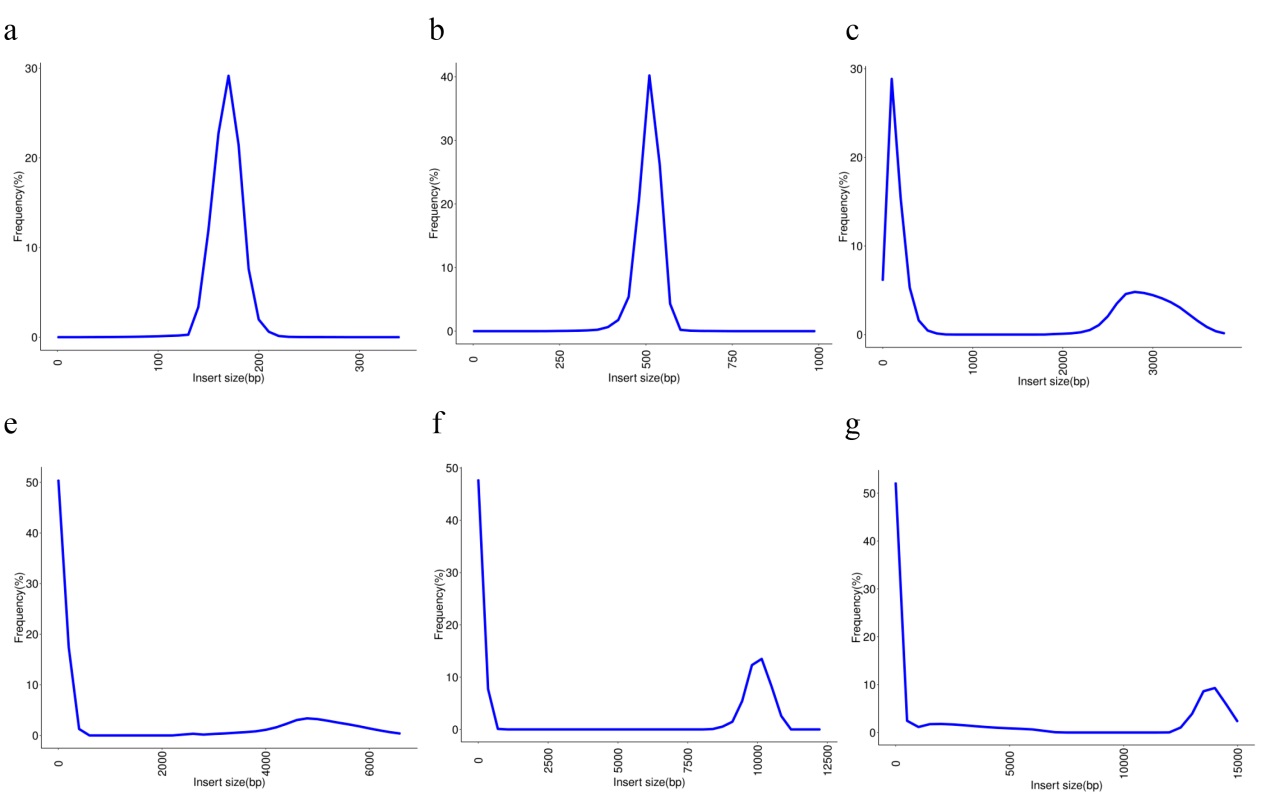

**Figure S3. Thirty-five sparidae specific genes involved in cellular processes, and environmental information processing in the *A. latus* genome.**

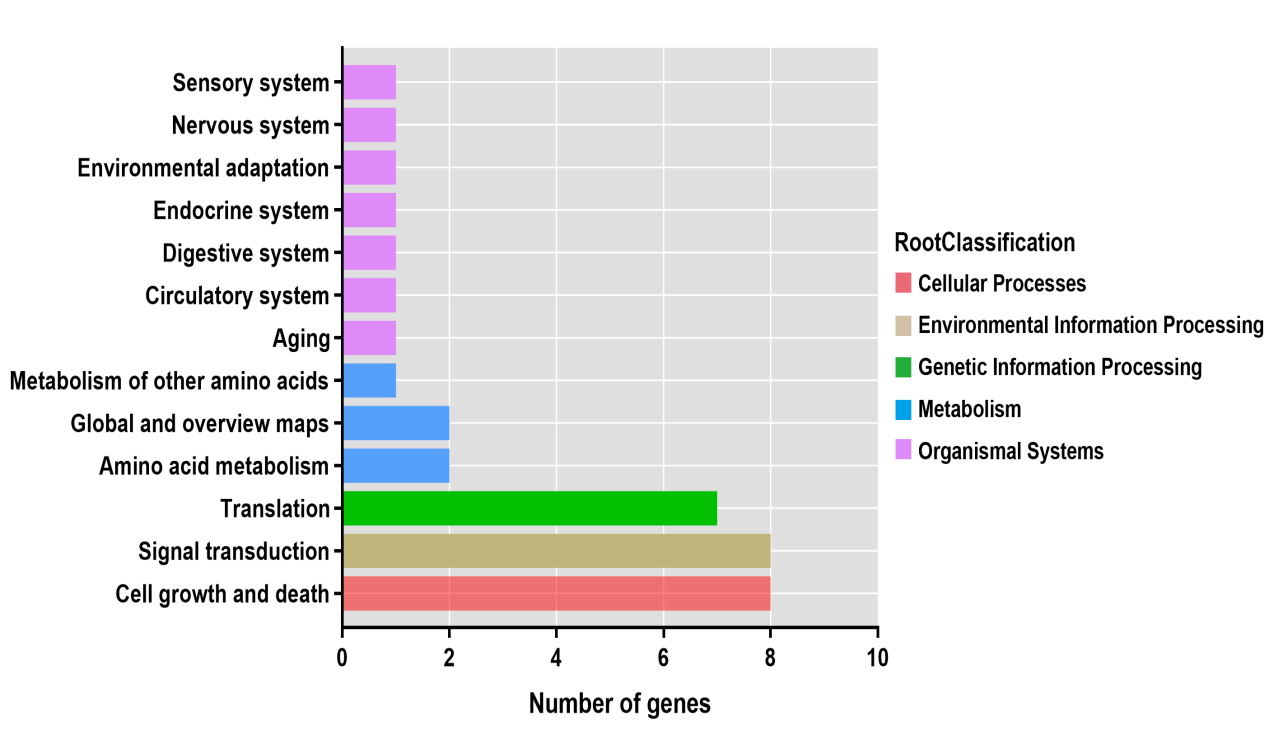

**Figure S4. Distribution of expanded genes in GO annotations indicative of abundance in integral component of membrane, ion binding and nucleic acid binding .**

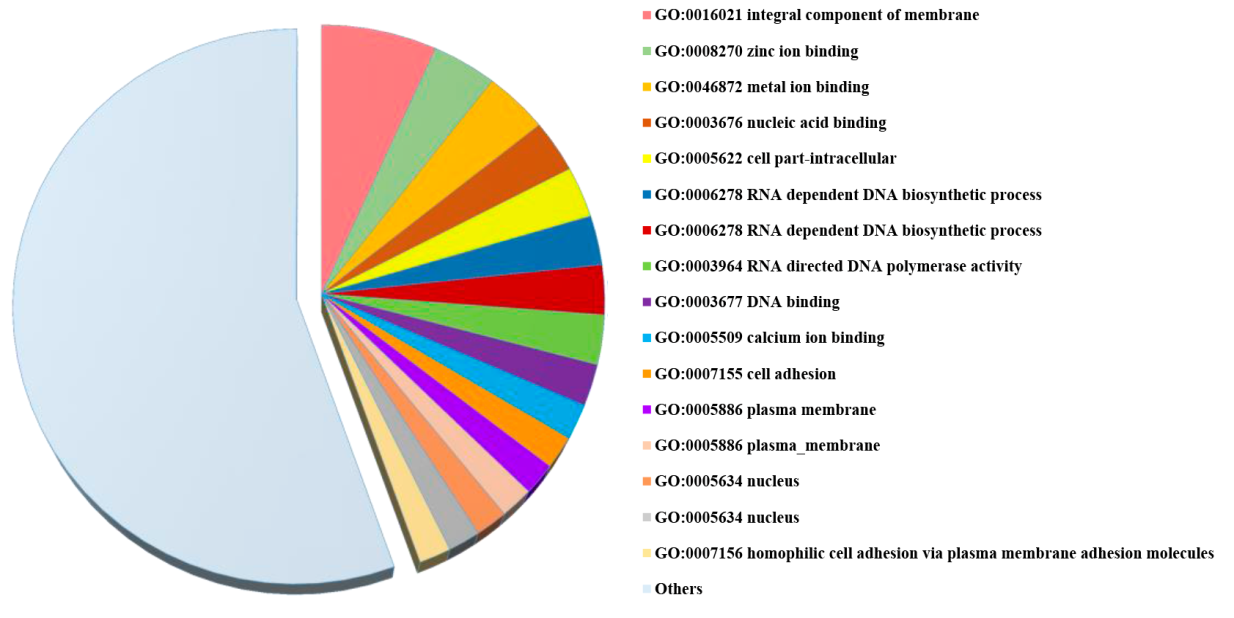

**Figure S5. Over-represented KEGG pathways for *A. latus* expanded gene families.**

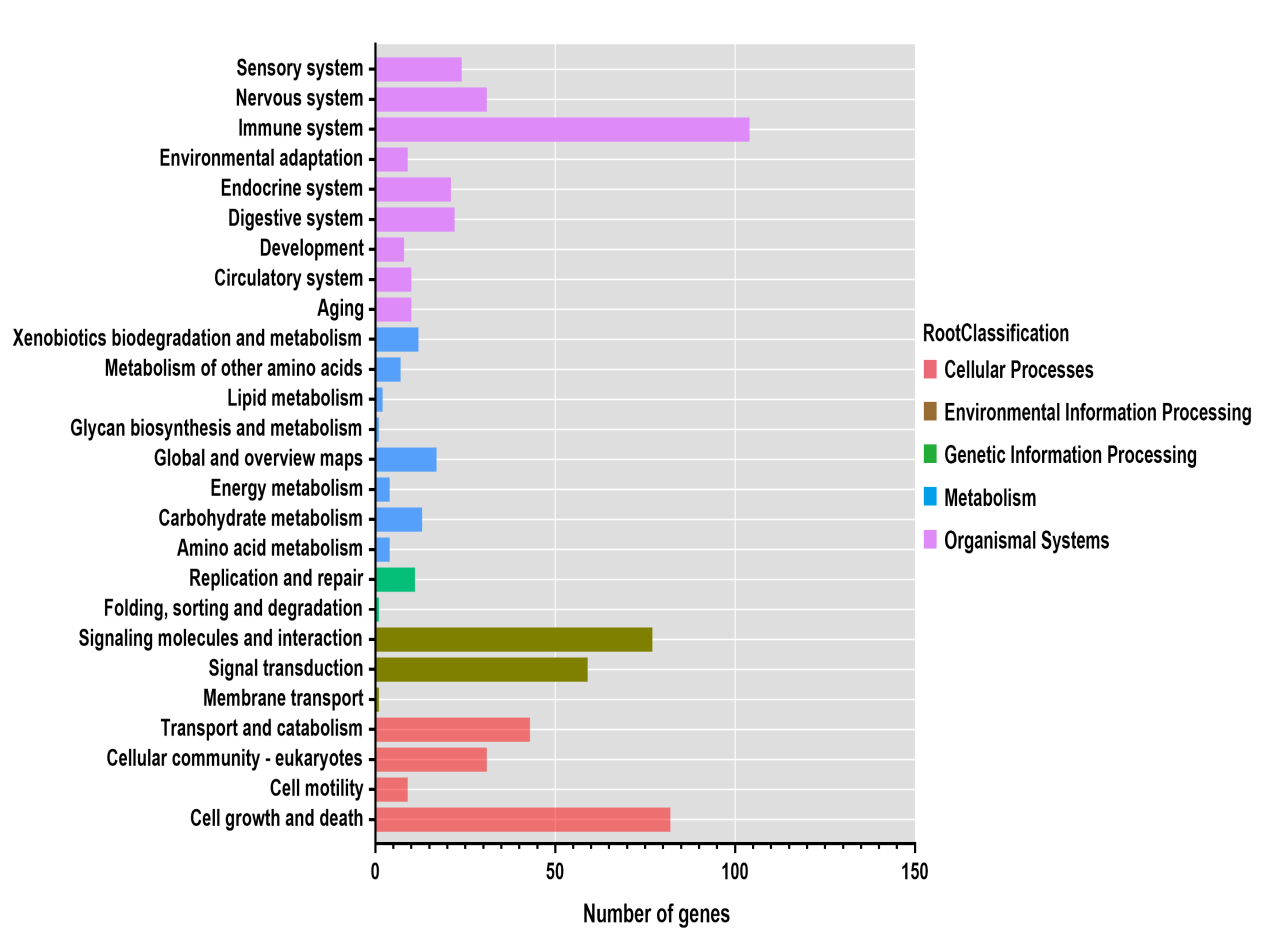

**Figure S6.** Phylogenetic tree of the *GSTA* gene family in teleosts. Three clades of *GSTA* in the *A. latus* genome were observed, the expanded genes are marked by red stars. The genes in largest clade are specifically expanded *GSTA* in the *A. latus* genome, which are also tandemly duplicated. The arrow indicates the transcriptional orientation.

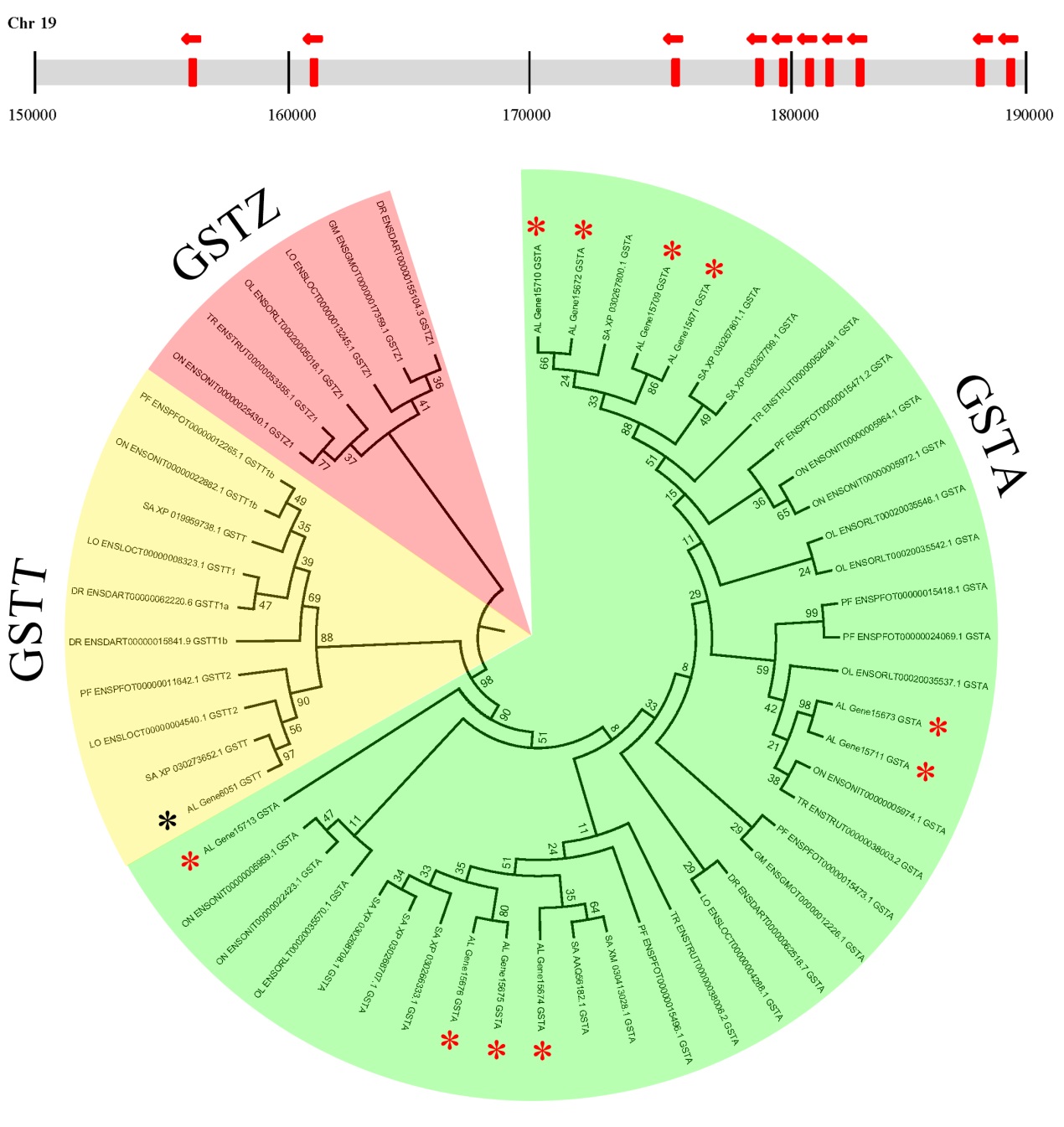

**Figure S7.** **Genome structure of the *GSTA* gene in teleosts.**

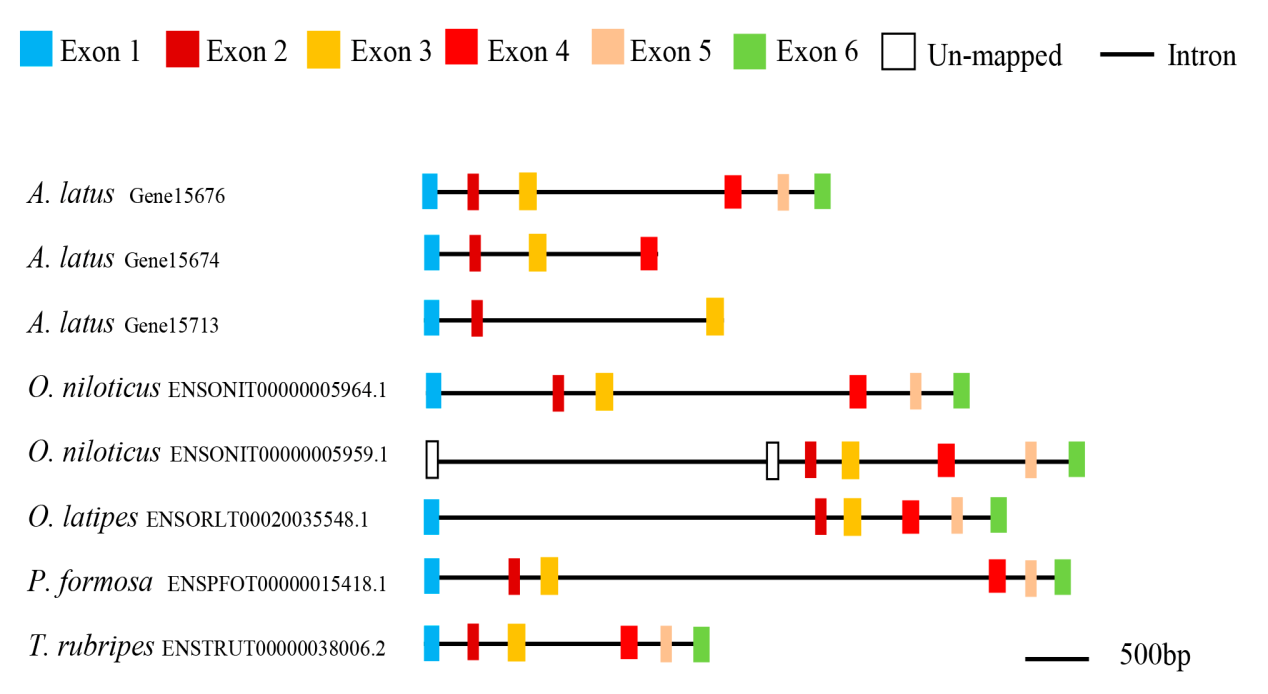

**Figure S8.** Temporal expression of four *GSTA* genes in gill, kidney, skin, and brain after acute salinity stress (low and high salinity) for the indicated time points.

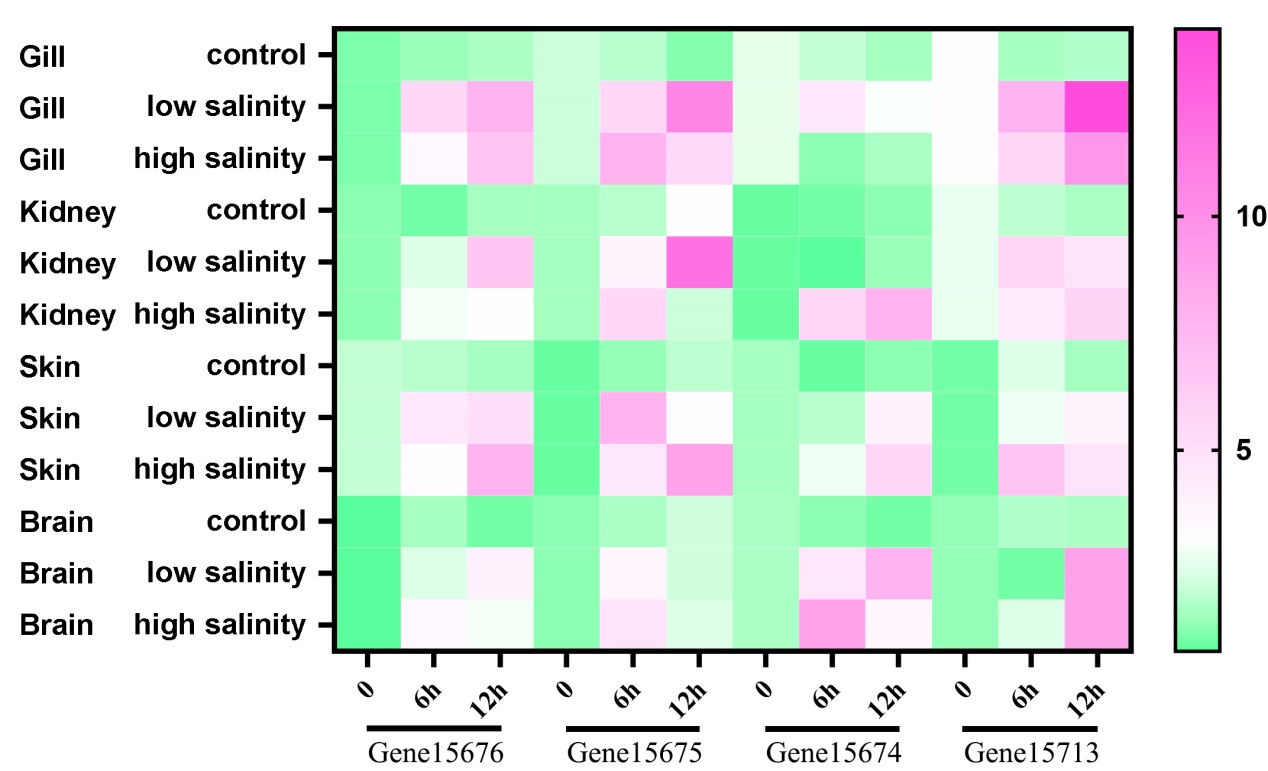

**Figure S9.** **Comparison of the expanded gene families, SLC12A2 (NKCC1) and** **SLC12A5. The areas of circles are proportional to the size of the gene family.**

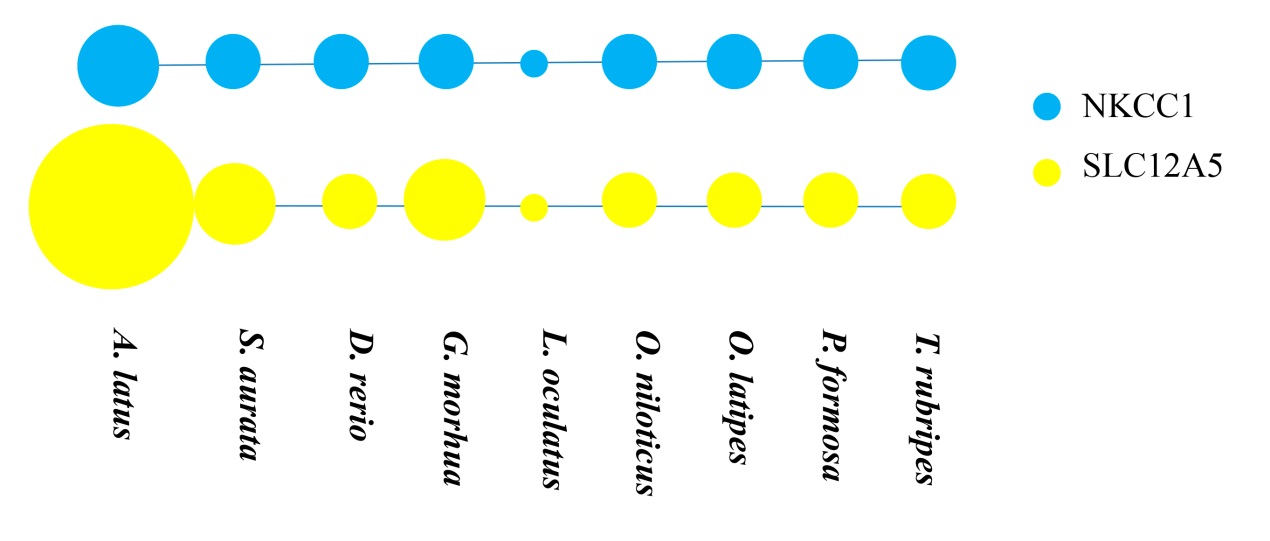

**Figure S10.** Temporal expression of four *SLC12A5* genes in gill, kidney, skin, and brain after acute salinity stress (low and high salinity) for the indicated time points.

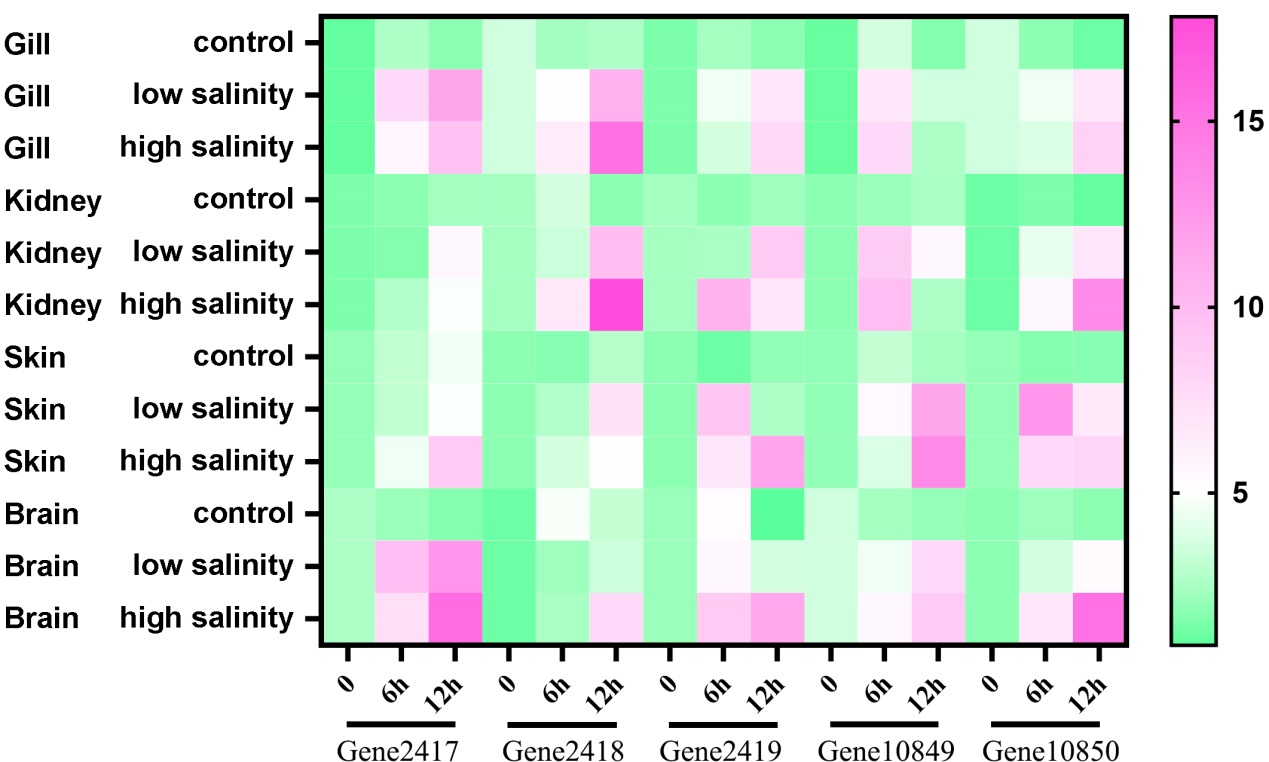

**Figure S11. Top 20 statistics of pathway enrichment for stage1-vs-stage2, stage1-vs-stage3.**

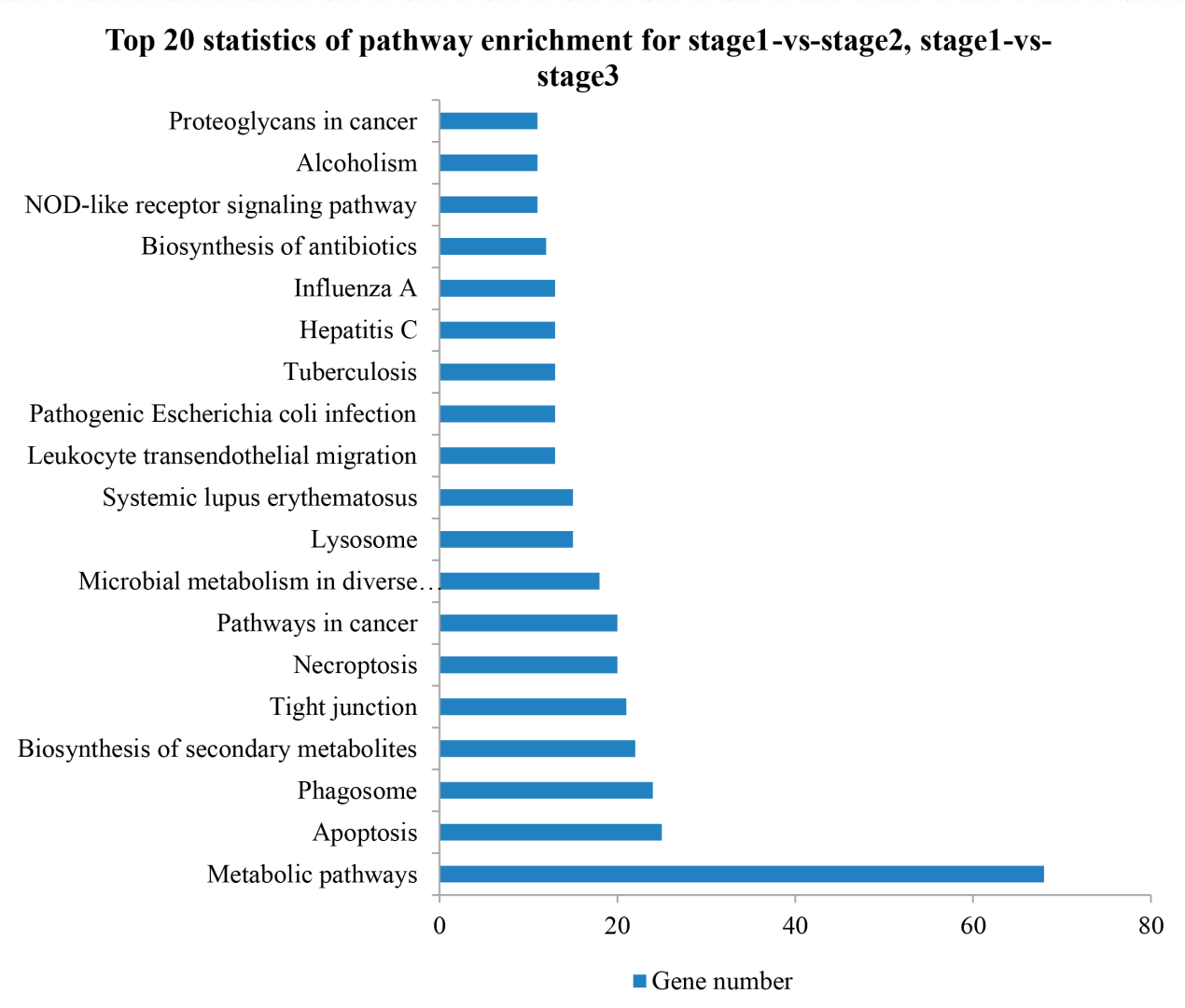

**Figure S12. Differential genes expression in the stage 3-vs-stage 2 (a), stage 3-vs-stage 1(b), and stage 2-vs-stage 1 (c) by RNA-seq and qRT-PCR.**

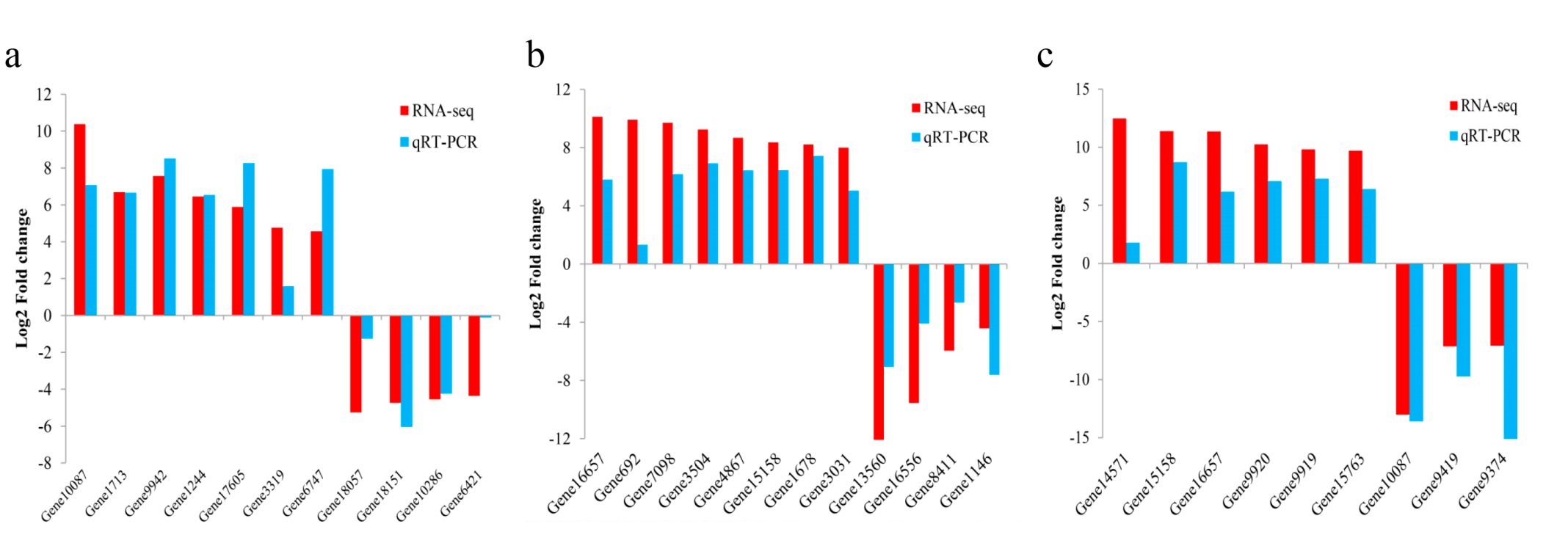

**Supplementary Table**

**Table S1. Mapping ratio of RNA-seq reads from three tissues in *A. latus***

| **SampleID** | **SEX** | **Pairs** | **Alignment Number** | **Alignment ratio** | **SRA Accesion Number** |
| --- | --- | --- | --- | --- | --- |
| R1703754_HCTKKCCXY | M1 | 36577847 | 33970510 | 92.87% | SRR10249786 |
| R1703755_HCTKKCCXY | M2 | 26075350 | 24353640 | 93.40% | SRR10249785 |
| R1703757_HCTKKCCXY | M3 | 36603034 | 33992006 | 92.87% | SRR10249794 |
| R1703759_HCTKKCCXY | In1 | 46675122 | 42715328 | 91.52% | SRR10249793 |
| R1703760_HCTKKCCXY | In2 | 26557817 | 24135639 | 90.88% | SRR10249792 |
| R1704084_HCTKKCCXY | In3 | 25545116 | 23271879 | 91.10% | SRR10249791 |
| R1703761_HCTKKCCXY | Fe1 | 27305258 | 25425184 | 93.11% | SRR10249790 |
| R1703762_HCTKKCCXY | Fe2 | 39493629 | 35996891 | 91.15% | SRR10249789 |
| R1704083_HCTKKCCXY | Fe3 | 48999037 | 44332893 | 90.48% | SRR10249788 |

**Table S2. BUSCO evaluation of the *A. latus* genes compared with the vertebrate gene set**

| **BUSCO benchmark** | **Number** | **Percentage (%)** |
| --- | --- | --- |
| Complete BUSCOs | 2385 | 92.23% |
| Complete and single-copy BUSCOs | 869 | 33.60% |
| Complete and duplicated BUSCOs | 1516 | 58.62% |
| Fragmented BUSCOs | 112 | 4.33% |
| Missing BUSCOs | 89 | 3.44% |
| Total BUSCO vertebrate genes | 2586 | 100.00% |

**Table S3. Detailed classification of repeat sequences in the assembled *A. latus* genome.**

|  | **Repbase TEs** | | **TE proteins** | | **Denovo TEs** | | **Combined TEs** | |
| --- | --- | --- | --- | --- | --- | --- | --- | --- |
| **DNA** | 584,175 | 0.08% | 704,579 | 0.10% | 9,865,645 | 1.35% | 11,044,136 | 1.51% |
| **LINEs** | 931,907 | 0.13% | 9,889,780 | 1.35% | 9,107,763 | 1.24% | 14,304,242 | 1.95% |
| **LTRs** | 220,596 | 0.03% | 6,678,283 | 0.91% | 1,944,340 | 0.27% | 7,341,300 | 1.00% |
| **SINE** | 83,845 | 0.01% | 0 | 0.00% | 0 | 0.00% | 84,508 | 0.01% |
| **Unclassified** | 6,088 | 0.01% | 0 | 0.00% | 106,163,538 | 14.50% | 106,163,538 | 14.50% |
| **Total** | 1,826,611 | 0.25% | 17,272,642 | 2.36% | 127,081,286 | 17.36% | 138,937,724 | 18.98% |

**Table S4. Detailed classification of repeat sequences in the assembled Sparus aurata genome.**

|  | **Repbase TEs** | | **TE proteins** | | **Denovo TEs** | | **Combined TEs** | |
| --- | --- | --- | --- | --- | --- | --- | --- | --- |
| **DNA** | 14,178,599 | 1.71% | 10,947,920 | 1.32% | 55,878,677 | 6.73% | 70,343,170 | 8.47% |
| **LINEs** | 10,682,809 | 1.29% | 21,791,455 | 2.62% | 12,981,643 | 1.56% | 31,476,885 | 3.79% |
| **LTRs** | 5,198,586 | 0.63% | 24,748,239 | 2.98% | 5,427,853 | 0.65% | 30,146,085 | 3.63% |
| **SINE** | 525,412 | 0.06% | 0 | 0.00% | 2,535,910 | 0.31% | 2,689,882 | 0.32% |
| **Unclassified** | 480,480 | 0.06% | 0 | 0.00% | 76,267,713 | 9.18% | 76,267,713 | 9.18% |
| **Total** | 31,065,886 | 3.74% | 57,487,614 | 6.92% | 153,091,796 | 18.44% | 210,923,735 | 25.40% |

**Table S5. Detailed classification of repeat sequences in the assembled *Acanthopagrus schlegelii* genome.**

|  | **Repbase TEs** | | **TE proteins** | | **Denovo TEs** | | **Combined TEs** | |
| --- | --- | --- | --- | --- | --- | --- | --- | --- |
| **DNA** | 20.930 | 3.041 % | 2.200 | 0.320% | 58.340 | 8.479 % | 68.130 | 9.902% |
| **LINEs** | 10.240 | 1.488 % | 6.950 | 1.010% | 26.760 | 3.889 % | 33.020 | 4.789 % |
| **LTRs** | 1.120 | 0.163 % | 2.340 | 0 | 3.780 | 0.550% | 4.550 | 0.661 % |
| **SINE** | 7.200 | 1.046 % | 35.410 | 0.340% | 25.980 | 3.062% | 31.270 | 4.544% |
| **Unclassified** | 0.020 | 0.003% | 0 | 0 | 0 | 0 | 0.020 | 0.003 % |
| **Total** | 0 | 0 | 0 | 0 | 25.370 | 3.687% | 25.370 | 3.687% |

**Table S6. Chromosome length and number of scaffolds from the assembled *A. latus* genome**

| **Chromosome No.** | **Length (bp)** | **Number of**  **scaffolds** |
| --- | --- | --- |
| LG1 | 37,811,058 | 206 |
| LG2 | 37,657,891 | 154 |
| LG3 | 37,224,566 | 182 |
| LG4 | 34,387,756 | 175 |
| LG5 | 33,952,217 | 144 |
| LG6 | 33,860,421 | 143 |
| LG7 | 33,758,209 | 141 |
| LG8 | 33,524,615 | 151 |
| LG9 | 33,500,224 | 142 |
| LG10 | 33,116,182 | 139 |
| LG11 | 32,993,337 | 158 |
| LG12 | 30,688,393 | 138 |
| LG13 | 30,257,479 | 117 |
| LG14 | 29,559,660 | 118 |
| LG15 | 28,614,909 | 123 |
| LG16 | 28,483,669 | 145 |
| LG17 | 28,241,308 | 144 |
| LG18 | 28,167,338 | 111 |
| LG19 | 27,743,381 | 149 |
| LG20 | 27,175,577 | 125 |
| LG21 | 26,291,649 | 97 |
| LG22 | 24,503,382 | 131 |
| LG23 | 23,194,532 | 102 |
| LG24 | 17,436,492 | 79 |
| Average | 30,506,010 | 206 |
| Total | 732,144,245 | 3314 |

**Table S7. Genomic collinear regions of *A. latus* and *L. crocea***

| **Chromosome** | **Link number** |
| --- | --- |
| LG1 | 542 |
| LG2 | 669 |
| LG3 | 671 |
| LG4 | 1059 |
| LG5 | 630 |
| LG6 | 691 |
| LG7 | 710 |
| LG8 | 603 |
| LG9 | 680 |
| LG10 | 579 |
| LG11 | 369 |
| LG12 | 528 |
| LG13 | 628 |
| LG14 | 378 |
| LG15 | 324 |
| LG16 | 579 |
| LG17 | 506 |
| LG18 | 399 |
| LG19 | 372 |
| LG20 | 435 |
| LG21 | 537 |
| LG22 | 458 |
| LG23 | 301 |
| LG24 | 167 |

**Table S8. Distribution of different orthologues genes from representative teleost genome. “1:1:1” indicates universal single-copy genes. “X:X:X” indicates orthologues exist in all representative genomes (missing in one species not allowed), with “X” meaning one or more orthologs per species. Those “1:1:1” genes were not included in “X:X:X”. “Sparidae specific” indicates genes only exist in the Sparidae fish. “Teleost specific” indicates orthologues exist in all teleost genomes. “Others” indicates genes which are assigned one gene family but do not fit into the categories of “1:1:1”, “X:X:X”, “Teleost specific”, and “species-specific”. “species-specific” includes genes without homologs to other species.**

| **Name** | **1:1:1** | **X:X:X** | **Sparidae specific** | **Species specific** | **Other** | **All** **assigned** | **Genome size(Mb)** |
| --- | --- | --- | --- | --- | --- | --- | --- |
| *Acanthopagrus latus* | 367 | 8122 | 35 | 2170 | 8937 | 19631 | 806 |
| *Sparus aurata* | 367 | 6647 | 31 | 7519 | 11380 | 25944 | 830 |
| *Acanthopagrus schlegelii* | 367 | 6862 | 25 | 539 | 11672 | 19465 | 688 |
| *Danio rerio* | 367 | 9667 | 0 | 2016 | 18263 | 30313 | 1674 |
| *Gadus morhua* | 367 | 7160 | 0 | 1103 | 11465 | 20095 | 608 |
| *Labrus bergylta* | 367 | 8778 | 0 | 1440 | 16808 | 27393 | 805 |
| *Lepisosteus oculatus* | 367 | 5756 | 0 | 795 | 11423 | 18341 | 869 |
| *Oreochromis niloticus* | 367 | 7923 | 0 | 318 | 12829 | 21437 | 816 |
| *Oryzias latipes* | 367 | 7647 | 0 | 1137 | 14471 | 23622 | 734 |
| *Poecilia formosa* | 367 | 8111 | 0 | 243 | 14894 | 23615 | 714 |
| *Takifugu rubripes* | 367 | 7171 | 0 | 600 | 12407 | 20545 | 391 |
| *Tetraodon nigroviridis* | 367 | 7396 | 0 | 596 | 11243 | 19602 | 342 |
| *Xiphophorus maculatus* | 367 | 7809 | 0 | 725 | 14873 | 23774 | 704 |
| *Cyprinus carpio* | 367 | 13636 | 0 | 5828 | 29446 | 49277 | 1714 |
| *Larimichthys crocea* | 367 | 7973 | 0 | 360 | 14667 | 23367 | 658 |
| *Notothenia coriiceps* | 367 | 7916 | 0 | 1909 | 14597 | 24789 | 637 |

**Table S9. Thirty-five sparidae specific genes involved in cellular processes, and environmental information processing in the *A. latus* genome.**

| **Gene ID** | **Gene description** | **Gene ID** | **Gene description** |
| --- | --- | --- | --- |
| Gene1087 | low affinity immunoglobulin gamma Fc region receptor III-like | Gene7916 | uncharacterized protein LOC109062443 |
| Gene1154 | uncharacterized protein | Gene7999 | Transposon TX1 hypothetical protein |
| Gene1228 | Transposon TX1 hypothetical protein | Gene8274 | uncharacterized protein |
| Gene1867 | Transposon TX1 hypothetical protein | Gene8276 | uncharacterized protein |
| Gene1874 | Transposon TX1 hypothetical protein | Gene9648 | hypothetical protein cypCar_00000299 |
| Gene3530 | putative transposase element L1Md-A101/L1Md-A102/L1Md-A2 | Gene11678 | uncharacterized protein LOC109062443 |
| Gene3642 | Transposon TX1 hypothetical protein | Gene12994 | cytosolic non-specific dipeptidase-like |
| Gene3745 | uncharacterized protein LOC106911834 isoform X1 | Gene14535 | uncharacterized protein LOC109062443 |
| Gene3756 | uncharacterized protein LOC103358552 | Gene15451 | 14-3-3 protein beta/alpha-like protein |
| Gene3893 | uncharacterized protein LOC109062443 | Gene15855 | Reverse transcriptase-like protein |
| Gene4143 | HBS1-like protein isoform X2 | Gene16030 | GTPase IMAP family member 8 |
| Gene4370 | Transposon TX1 hypothetical protein | Gene16608 | Reverse transcriptase-like protein |
| Gene5156 | Transposon TX1 hypothetical protein | Gene16631 | Transposon TX1 hypothetical protein |
| Gene5244 | uncharacterized protein LOC108281047 | Gene17890 | Phosphatase and actin regulator 1 |
| Gene5451 | neurofilament heavy polypeptide | Gene17908 | Phosphatase and actin regulator 1 |
| Gene5457 | neurofilament heavy polypeptide | Gene19569 | LINE-1 type transposase domain-containing protein 1 |
| Gene6053 | uncharacterized protein LOC109062443 | Gene19573 | transmembrane emp24 domain-containing protein 9-like |
| Gene7297 | Paternally-expressed 3 protein |  |  |

**Table S10. Expansion and contraction of gene families (gene numbers <100) among nine teleost species.**

| **Name** | **Expansions** | **Genes Gained** | **Equal** | **Contractions** | **Genes Lost** | **Families Lost** | **Average Expansion** | **Sig Expansions** | **Sig Contractions** | **Total Sig Changes** |
| --- | --- | --- | --- | --- | --- | --- | --- | --- | --- | --- |
| *Acanthopagrus latus* | 1620 | 3567 | 12334 | 5206 | 5856 | 4799 | -0.11947 | 238 | 87 | 325 |
| *Sparus aurata* | 1465 | 2842 | 14006 | 3689 | 4204 | 2859 | -0.07109 | 271 | 98 | 369 |
| *Takifugu rubripes* | 835 | 1168 | 15037 | 3288 | 3532 | 2506 | -0.12338 | 36 | 37 | 73 |
| *Poecilia formosa* | 1577 | 2648 | 16470 | 1113 | 1173 | 850 | 0.076983 | 145 | 6 | 151 |
| *Oryzias latipes* | 828 | 2358 | 16709 | 1623 | 1802 | 1139 | 0.029019 | 163 | 27 | 190 |
| *Oreochromis niloticus* | 813 | 1938 | 15743 | 2604 | 2714 | 2247 | -0.04050 | 111 | 11 | 122 |
| *Gadus morhua* | 730 | 1110 | 13351 | 5079 | 5339 | 4179 | -0.22072 | 21 | 19 | 40 |
| *Danio rerio* | 4456 | 8085 | 10470 | 4234 | 4357 | 3483 | 0.194572 | 88 | 3 | 91 |
| *Lepisosteus oculatus* | 704 | 1466 | 11158 | 7298 | 7837 | 4811 | -0.33252 | 28 | 12 | 40 |

**Table S11. Expansion and contraction of gene families (gene numbers >100) among nine teleost species.**

| **Name** | **Expansions** | **Genes Gained** | **Equal** | **Contractions** | **Genes Lost** | **Families Lost** | **Average Expansion** | **Sig Expansions** | **Sig Contractions** | **Total Sig Changes** |
| --- | --- | --- | --- | --- | --- | --- | --- | --- | --- | --- |
| *Acanthopagrus latus* | 7 | 852 | 6 | 2 | 17 | 1 | 55.6667 | 0 | 0 | 0 |
| *Sparus aurata* | 1 | 109 | 9 | 5 | 75 | 3 | 2.26667 | 0 | 0 | 0 |
| *Takifugu rubripes* | 0 | 0 | 6 | 9 | 37 | 7 | -2.46667 | 0 | 0 | 0 |
| *Poecilia formosa* | 1 | 3 | 9 | 5 | 16 | 3 | -0.866667 | 0 | 0 | 0 |
| *Oryzias latipes* | 3 | 200 | 9 | 3 | 3 | 2 | 13.1333 | 0 | 0 | 0 |
| *Oreochromis niloticus* | 2 | 84 | 7 | 6 | 10 | 5 | 4.93333 | 0 | 0 | 0 |
| *Gadus morhua* | 2 | 233 | 1 | 12 | 28 | 10 | 13.6667 | 0 | 0 | 0 |
| *Danio rerio* | 5 | 738 | 1 | 9 | 18 | 5 | 48 | 0 | 0 | 0 |
| *Lepisosteus oculatus* | 0 | 0 | 1 | 14 | 89 | 12 | -5.93333 | 0 | 0 | 0 |
